# Derivation of Airway Basal Stem Cells from Human Pluripotent Stem Cells

**DOI:** 10.1101/2020.02.21.959395

**Authors:** Finn J. Hawkins, Shingo Suzuki, Mary Lou Beermann, Cristina Barillà, Ruobing Wang, Carlos Villacorta-Martin, Andrew Berical, J.C. Jean, Jake Le Suer, Chantelle Simone-Roach, Yang Tang, Thorsten M. Schlaeger, Ana M. Crane, Sarah X. L. Huang, Scott H. Randell, Andras Rab, Eric J. Sorscher, Amjad Horani, Steven L. Brody, Brian R. Davis, Darrell N. Kotton

## Abstract

The derivation of self-renewing tissue-specific stem cells from human induced pluripotent stem cells (iPSCs) would shorten the time needed to engineer mature cell types *in vitro* and would have broad reaching implications for the field of regenerative medicine. Here we report the directed differentiation of human iPSCs into putative airway basal cells (“iBCs”), a population resembling the epithelial stem cell of lung airways. Using a dual fluorescent reporter system (NKX2-1^GFP^;TP63^tdTomato^) we track and purify these cells over time, as they first emerge from iPSC-derived foregut endoderm as developmentally immature NKX2-1^GFP+^ lung progenitors which then augment a TP63 program during subsequent proximal airway epithelial patterning. These cells clonally proliferate, initially as NKX2-1^GFP+^/TP63^tdTomato+^ immature airway progenitors that lack expression of the adult basal cell surface marker NGFR. However, in response to primary basal cell medium, NKX2-1^GFP+^/ TP63^tdTomato+^ cells upregulate NGFR and display the molecular and functional phenotype of airway basal stem cells, including the capacity to clonally self-renew or undergo multilineage ciliated and secretory epithelial differentiation in air-liquid interface cultures. iBCs and their differentiated progeny recapitulate several fundamental physiologic features of normal primary airway epithelial cells and model perturbations that characterize acquired and genetic airway diseases. In an asthma model of mucus metaplasia, the inflammatory cytokine IL-13 induced an increase in MUC5AC+ cells similar to primary cells. CFTR-dependent chloride flux in airway epithelium generated from cystic fibrosis iBCs or their syngeneic CFTR-corrected controls exhibited a pattern consistent with the flux measured in primary diseased and normal human airway epithelium, respectively. Finally, multiciliated cells generated from an individual with primary ciliary dyskinesia recapitulated the ciliary beat and ultrastructural defects observed in the donor. Thus, we demonstrate the successful *de novo* generation of a tissue-resident stem cell-like population *in vitro* from iPSCs, an approach which should facilitate disease modeling and future regenerative therapies for a variety of diseases affecting the lung airways.

## Introduction

Basal cells (BCs) of the adult mouse and human airways are capable of self-renewal and multi-lineage differentiation under homeostatic conditions *in vivo*, in response to injury, and after culture expansion *ex vivo*, thereby fulfilling the definition of a tissue-specific adult stem cell (Rock et al. 2009). An extensive literature has established that BCs can regenerate the airway epithelium by serving as precursors for essential specialized epithelial cell-types including secretory and multiciliated cells (Rock et al. 2009; Montoro et al. 2018; Plasschaert et al. 2018). These stem cell properties make BCs a highly desirable cell type to generate *ex vivo* for modeling airway diseases and a leading candidate for cell-based therapies designed to reconstitute the airway epithelium. However, to date, the *de novo* derivation in culture of bona fide self-renewing basal cells that are similar to their primary cell counterparts has yet to be achieved, although several recent reports have demonstrated the successful differentiation of human pluripotent stem cells into airway epithelial cell types, including those that express the canonical basal cell marker TP63 (Hawkins et al. 2017; McCauley et al. 2017; Konishi et al. 2016; Dye et al. 2015). The derivation of self-renewing tissue-specific stem cells, such as BCs from human induced pluripotent stem cells (iPSCs) would represent an important advance by shortening the time needed to engineer mature cell types *in vitro* and would have far reaching implications for the field of regenerative medicine.

Successful differentiation of iPSCs into BCs, as with other mature lineage targets, might be achieved if a detailed understanding of the embryonic development and mature adult phenotype of the desired lineage were used as a guide to *in vitro* culture optimization. In humans, BCs are located in many tissues including the airway, nasal, olfactory, urinary tract, skin, and esophageal epithelium. In human airways, BCs are highly abundant in the pseudostratified epithelium extending from the trachea to the terminal bronchioles (Rock et al. 2010). Airway BCs can be identified based on their classic anatomic location along the basal lamina and by the expression of several markers including Tumor Protein 63 (TP63), cytoskeletal protein Keratin 5 (KRT5) and Nerve Growth Factor Receptor (NGFR) (Rock et al. 2010). TP63, a member of the p53 family of transcription factors, is essential to the BC program in the airway but also other organs (A. Yang et al. 1999).

While airway basal cells in adult lungs have been extensively studied for many years, only recently has their developmental origin been examined. For example, lineage-tracing experiments in mice (Y. Yang et al. 2018) reveal that a Tp63-program is already present early in lung development at the time of initial lung bud formation (embryonic day E9.5) within a subset of lung epithelial progenitors expressing the transcriptional regulator that marks all developing lung epithelial cells, NK2 homeobox 1 (Nkx2-1). The early Nkx2-1+/Tp63+ co-expressing cells are not basal cells since they lack the BC morphology and molecular program. Rather these fetal cells function as multipotent progenitors of subsequent alveolar and airway epithelia (Y. Yang et al. 2018). Tp63 expression is then gradually restricted to the developing airways where it is initially broadly expressed in immature airway progenitors and later restricted to a subset of tracheal cells that localize to the basement membrane and upregulate markers of adult BCs, including Krt5 and Ngfr. A rare population of Tp63+/Krt5-cells, thought to be immature BCs, persists in the intrapulmonary airways of adult mice (Y. Yang et al. 2018). The signaling pathways that control basal cell specification and maturation in the lung are not precisely known; however, in-bred mouse models suggest a temporal role for FGF10/FGFR2b (Volckaert et al. 2013). Although limited data are available regarding the developmental origins of BCs in humans, a similar pattern to that observed in mice of early and diffuse TP63 expression in the airway epithelium followed by restricted expression in KRT5+ BCs has also been described (Nikolić et al. 2017). That said, extrapolating findings from mouse BCs to humans is challenging since the location of BCs in adult humans (throughout the conducting airways) differs markedly from mice, where mature BCs localize predominantly to the trachea and large extra-pulmonary airways.

Given the stem cell properties of airway BCs including their established proliferative capacity, well-established protocols have been developed to expand primary human BCs *in vitro* (Fulcher & Randell 2013; Mou et al. 2016; Suprynowicz et al. 2017). These BCs, conventionally referred to as human bronchial epithelial cells (HBECs), differentiate into a pseudostratified, multicellular, airway epithelium in air-liquid interface (ALI) culture that recapitulates aspects of *in vivo* airway biology. This approach has been used to study airway progenitor populations, cell fate decisions, and plasticity within the airway epithelium (Rock et al. 2011; Plasschaert et al. 2018). The understanding of acquired and genetic human airway diseases, including the mucus metaplasia of asthma, the chloride transport defects of cystic fibrosis, and the ciliary dysfunction of primary ciliary dyskinesia, has advanced through this model (Seibold 2018; Clancy et al. 2019; Horani et al. 2016).

Despite their utility, HBECs as a source of primary basal cells also have significant shortcomings that limit their application for lung research. HBECs are isolated in small quantities from airway brushings during bronchoscopy or in larger quantities from either lung transplant recipient explant tissue or donor organs that are not used for transplant. As a result, there is a limited and unpredictable source of cells from patients with rarer genetic mutations such as nonsense *CFTR* variants or primary ciliary dyskinesia mutations. Furthermore, HBEC preparations from these individuals are often compromised by the secondary effects of infections, drugs, or ventilator injuries. In addition, as a potential source for cell-based airway therapies, approaches to generate sufficient primary BCs for airway engraftment would likely require an allogeneic source of cells thus requiring long term immunosuppression (Berical et al. 2019). Although successful *in vivo* engraftment of exogenous BCs is without precedent, iPSCs have the potential to overcome many of these hurdles: they are routinely generated from any patient while retaining the individual’s unique genetic background; they can be expanded in large numbers; and they are amenable to gene-editing and correction of disease-causing mutations. Thus, iPSCs can theoretically provide a source of autologous cells for future cell-based therapies.

While significant progress has been with regard to directed differentiation of iPSCs into a wide range of cell types, the approach is often limited by lengthy and complex differentiation protocols that result in heterogeneous cell types and limited numbers of the cell type of interest. Several groups, including ours, have reported the derivation of airway epithelial cells through directed differentiation of iPSCs (McCauley et al. 2017; Firth et al. 2014; Dye et al. 2015; Konishi et al. 2016; Huang et al. 2014). For example, we found that foregut endoderm derived from iPSCs can be specified into NKX2-1+ primordial lung epithelial progenitors through activation of Wnt, BMP, and RA signals (Serra et al. 2017) and then further patterned into proximal airway epithelium through downregulation of Wnt signals in an “airway medium” containing FGF2 and FGF10 (McCauley et al. 2017). These cultures contain cells with some markers found in BCs; however, the successful generation of bone-fide BCs with detailed characterization and demonstration of stable expansion and multi-lineage differentiation that are comparable to adult BCs has yet to be reported.

Here we successfully differentiate iPSCs *in vitro* into putative basal cells that share transcriptional similarities and key stem cell properties with their *in vivo* tissue-resident counterparts. The resulting approach recapitulates the sequence of key developmental milestones observed in mouse and human fetal lungs. Initially primordial lung progenitors identified by NKX2-1 expression are produced with only low levels of TP63 expression detectable in a minority of cells. Subsequently, an efficient developing airway program is induced characterized by co-expression of NKX2-1, TP63, and a key transcription factor in airway development, SOX2, with subsequent maturation into cells expressing the functional and molecular phenotype of BCs, including expression of the cell surface marker NGFR, that enables their purification by flow cytometry. The resulting sorted cells display long-term, clonal self-renewal capacity, trilineage differentiation in air-liquid interface (ALI) cultures *in vitro*, and can be applied for disease modeling studies, exemplified here by recapitulating the chloride channel dysfunction of CF, structural and motility defects of primary ciliary dyskinesia, and goblet cell hyperplasia of asthma. The ability to derive tissue-specific stem cells such as airway BCs from iPSCs, including capabilities to purify, expand, cryopreserve and differentiate these cells, overcomes many of the key hurdles that currently limit the more widespread application of iPSC technology and should accelerate the study of human lung disease.

## Results

### A TP63 Fluorescent Reporter Allows Visualization, Purification and Interrogation of iPSC-derived Airway Progenitors

Since NKX2-1 is the earliest transcriptional regulator expressed in all developing lung epithelial cells and TP63 is a canonical transcription factor required for expression of the basal cell program, we sought to generate a tool that would allow visualization and purification of NKX2-1+/TP63+ putative lung basal cells engineered *in vitro* from iPSCs while excluding any non-lung basal cells (NKX2-1-/TP63+)(Figure 1A-B). Using our published, normal iPSC line (BU3) that carries a GFP reporter targeted to the endogenous *NKX2-1* locus (Hawkins et al. 2017), we employed CRISPR/Cas9 gene editing to target a tdTomato reporter coding sequence into one allele of the endogenous *TP63* locus at exon 4 (Figure 1B). This strategy of targeting a reporter to exon 4 was taken with the goal of ensuring reporter expression could be visualized in the presence of expression of the predominant known forms of TP63, DeltaN-type or TA-type at the N-terminus while avoiding bias to C-terminal splice forms, such as α, β, and γ-type (Levrero et al. 2000). Karyotypically normal, pluripotent clones with monoallelic targeting of the *TP63* locus and biallelic targeting of the *NKX2-1* locus were identified (Figure S1A-E), thus establishing a biofluorescent reporter iPSC line, BU3 NKX2-1^GFP^;P63^tdTomato^, hereafter “BU3 NGPT”. To test faithfulness and specificity of the reporters, we differentiated this iPSC line into lung epithelium employing our recently published airway directed differentiation protocol in which we previously identified a population of basal-like cells based on the expression of NKX2-1, TP63 and KRT5 (Figure 1C) (McCauley et al. 2017). As expected, NKX2-1^GFP+^ (hereafter “GFP+”) cells, a small fraction of which also expressed tdTomato (hereafter “TOM+”), emerged by day 15 of differentiation as detected by flow cytometry and fluorescence microscopy (18.1±19% GFP+, 0.4±0.8% TOM+, mean±SD, 45.4% GFP+, 2.5% GFP+/TOM+ in the experiment shown in Figure 1E) (Figure 1D-E). Almost all TOM+ cells identified at this time point expressed the lung epithelial lineage GFP reporter (Figure 1E). To induce proximal airway differentiation of lung progenitors, GFP+ lung progenitor cells (regardless of TOM expression) were sorted on day 15 of the protocol and suspended in droplets of Matrigel in our previously published “airway” serum-free medium conditions containing FGF2, FGF10, dexamethasone, cyclic AMP, 3-isobutyl-1-methylxanthine and Y-27632, hereafter “FGF2+10+DCI+Y” (Figure 1C schematic) (McCauley et al. 2017). Between days 28-36, monolayered epithelial spheres emerged and 43.7±8.6% (mean±SD, 46.6% GFP+, 24.6% GFP+/TOM+ in the experiment shown in Figure 1E) of cells co-expressed NKX2-1^GFP^ and TP63^TOM^ (Figure 1E-F). With further single-cell dissociation of epithelial spheres and expansion in 3D culture conditions the percentage of cells co-expressing GFP and TOM, hereafter “GFP+/TOM+”, increased further to 73.7±4.7% by day 42 (Figure 1F). Flow cytometry sorting on day 36 demonstrated enrichment of *TP63* mRNA and protein in GFP+/TOM+ cells and depletion in TOM-cells (Figure 1G, H). TP63 protein was observed in few, if any cells that lacked TOM staining (Figure 1H). These findings confirmed specificity of the tdTomato reporter. In summary, we determined that NKX2-1^GFP+^/TP63^TOM+^ cells emerge and can be expanded in FGF2+10+DCI+Y medium. The early emergence of tdTomato expression in some cells beginning around day 15 is in keeping with our prior observations *in vitro* (Hawkins et al. 2017; McCauley et al. 2017; McCauley et al. 2018) and *in vivo* (Y. Yang et al. 2018; Nikolić et al. 2017) that *Tp63* mRNA and nuclear protein are initially detected at this time point in a subset of early NKX2-1+ primordial lung epithelial progenitors soon after specification of the respiratory lineage even prior to airway differentiation (see schematic Figure 1C).

**Figure 1:**
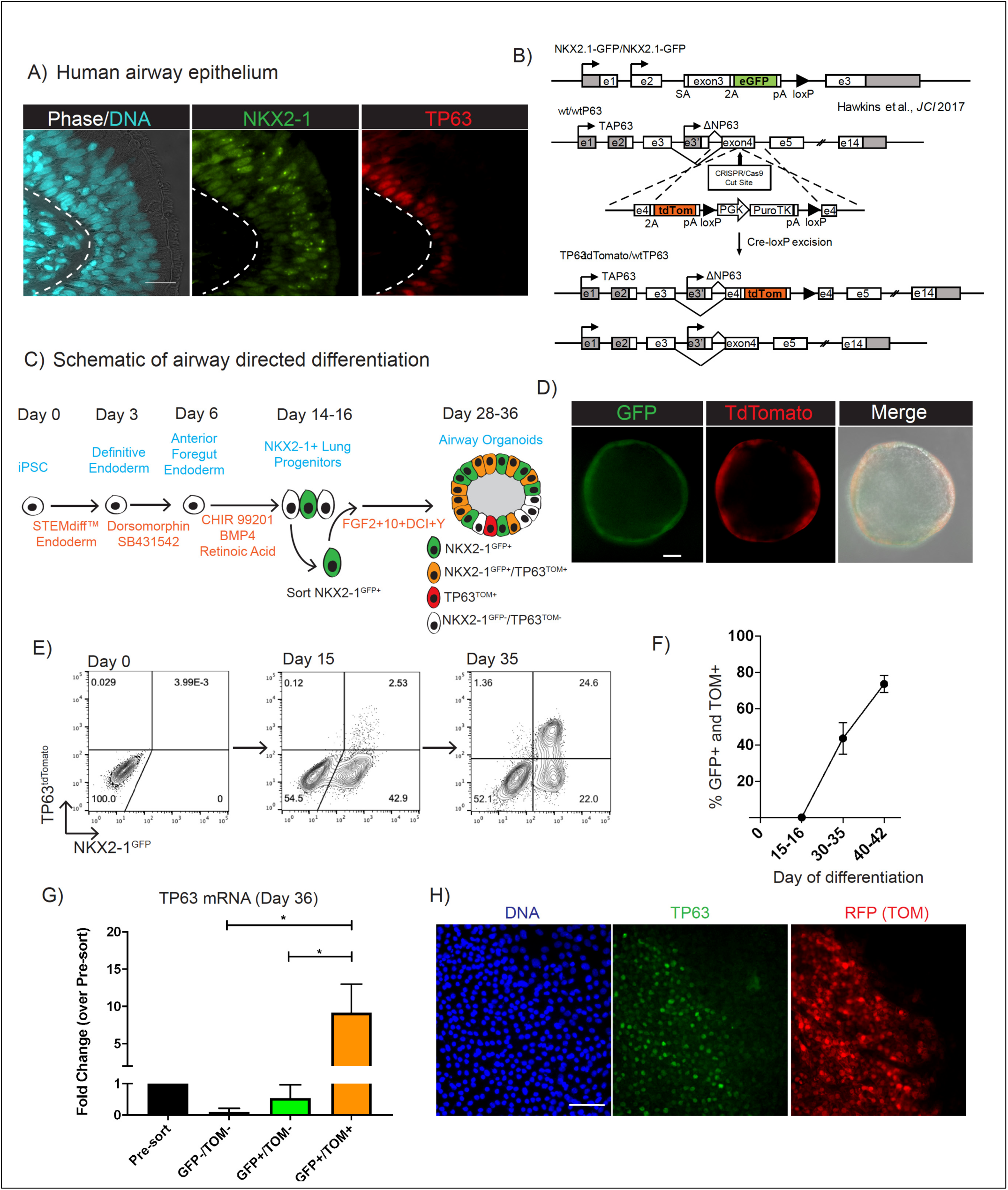
Generation of a dual fluorescent NKX2-1^GFP^;TP63^tdTomato^ iPSC reporter to track and purify putative basal cells. (A) Adult human airway immunolabeled with antibodies against NKX2-1 and TP63 (nuclei stained with Hoechst; scale bar=10μm). (B) Schematic of gene-editing strategy to insert tdTomato sequence into one allele of the *TP63* locus of previously generated NKX2-1^GFP^ iPSCs. See supplemental figure 1 for further details. (C) Schematic of airway directed differentiation protocol. Single positive NKX2-1^GFP+^ cells are indicated in green, TP63^tdTomato+^ in red and co-expression of NKX2-1^GFP+^/TP63^tdTomato+^ in orange. (D) Representative image of BU3 NGPT spheroid on day 36 of differentiation demonstrating GFP and tdTomato fluorescence (scale bar=50μm). (E) Representative flow cytometry plots of NKX2-1^GFP^ vs TP63^tdTomato^ expression on days 9, 15 and 35 of directed differentiation. (F) Quantification of the percentage of cells, calculated by flow cytometry, co-expressing NKX2-1^GFP^ and TP63^tdTomato^ between days 15-16 and 40-42 of differentiation (left panel, n=6). (G) qRT-PCR quantification of *TP63* mRNA levels (2^-ΔΔCt^) in GFP+/TOM+, GFP-/TOM-, and GFP+/TOM- populations sorted by FACS compared to presort levels on Day 36 of directed differentiation. (H) Immunolabeling of BU3 NGPT with antibodies against TP63 and RFP on day 30 of directed differentiation. The cells remained in 2D culture in FGF2+FGF10+DCI+Y medium from day 15 to facilitate antibody labeling (nuclei stained with DAPI; scale bar=50μm).

### iPSC-derived Airway Progenitors Adopt a Molecular Phenotype Similar to Primary Basal cells

We next sought to characterize NKX2-1^GFP+^/TP63^TOM+^ cells produced by the airway differentiation protocol in terms of capacity for self-renewal, multi-lineage differentiation, and expression of the canonical BC marker NGFR (Figures S2A-E). We sorted day 40-42 iPSC-derived GFP+/TOM+ cells and placed them in air-liquid interface (ALI) 2D cultures for at least 2 weeks and observed differentiation into cells expressing markers of multiciliated and secretory cells, although this differentiation appeared to be patchy (Figure S2A). iPSC-derived ALI cultures frequently failed to maintain barrier function evident as transepithelial electrical resistance (TEER) measurements (Figures S2A-B). In contrast, ALI cultures from primary HBECs differentiated into a consistent mucociliary epithelium that maintained barrier function and had significantly higher TEER measurements to iPSC-derived (Figures S2A-B). In keeping with these results, NGFR was expressed on only a small fraction, 1.9±1.8%, of GFP+/TOM+ cells on day 40-42 of differentiation, suggesting cells at this time point were unlikely to be mature basal cells. Even after 3 additional months of serial sphere passaging of GFP+/TOM+ cells in 3D culture, NGFR expression did not significantly increase, although GFP and TOM co-expression was maintained in the majority of cells (Figures S2C-E) suggesting that increased culture time alone would not induce BC maturation of iPSC-derived NKX2-1+/TP63+ cells. We concluded that in the presence of FGF2+10+DCI+Y, an early basal cell transcriptomic program is initiated (including co-expression of NKX2-1 and TP63), however, a key basal cell marker was lacking and these cells differentiated inconsistently in ALI culture (Figures S2A-E).

We reasoned that one possible explanation for the low expression of NGFR and inconsistent multi-lineage differentiation was that GFP+/TOM+ cells were more similar to immature TP63+/NKX2-1+/KRT5-airway progenitors observed during early airway patterning and were not yet mature basal cells (Y. Yang et al. 2018). Consistent with this speculation, KRT5 expression was detectable but low and variable in iPSC-derived GFP+/TOM+ cells compared to primary controls (Data not shown). Thus, we next sought to identify culture conditions that would further differentiate GFP+/TOM+ cells into a more basal-like state, using NGFR expression as a readout. Recently, inhibition of SMAD and ROCK signaling was found to induce a proliferative state in primary BCs and significantly increase overall yield from serially passaged cultures while maintaining, to an extent, their differentiation capacity (Mou et al. 2016; Zhang et al. 2018). We tested whether GFP+/TOM+ iPSC-derived airway progenitors would selectively proliferate and possibly mature in media conditions optimized for primary human BC culture. On day 30-32 of differentiation GFP+/TOM+ were sorted, replated in 3D Matrigel, and exposed to a commercially available BC medium with small molecules inhibitors of SMAD signaling (TGFβ inhibition and BMP inhibition with A 83-01 and DMH1, respectively) (Figures 2A-D). In response to PneumaCult-Ex Plus medium supplemented with SMAD inhibitors and Y-27632 (hereafter “BC medium”), GFP+/TOM+ spheroids where morphologically different based on their smaller, denser appearance and less distinct lumens (Figure 2A). We observed rapid and robust induction of NGFR in GFP+/TOM+ cells within 4 days of exposure to BC medium (Figures 2B-D and S2F-G). For example, between day 1 and day 3, NGFR expression increased in frequency from ∼1% to 35% in response to BC medium. By day 6, 71±5% (mean±SD) of cells expressed NGFR (Figure 2B), suggesting that the change in medium formulation, rather than selective expansion of rare NGFR+ cells, was inducing NGFR expression in previously NGFR-cells. In contrast, parallel aliquots of cells that continued in airway medium (FGF2+10+DCI+Y; Figure 2B, D, and S2F) expressed little to no NGFR.

**Figure 2:**
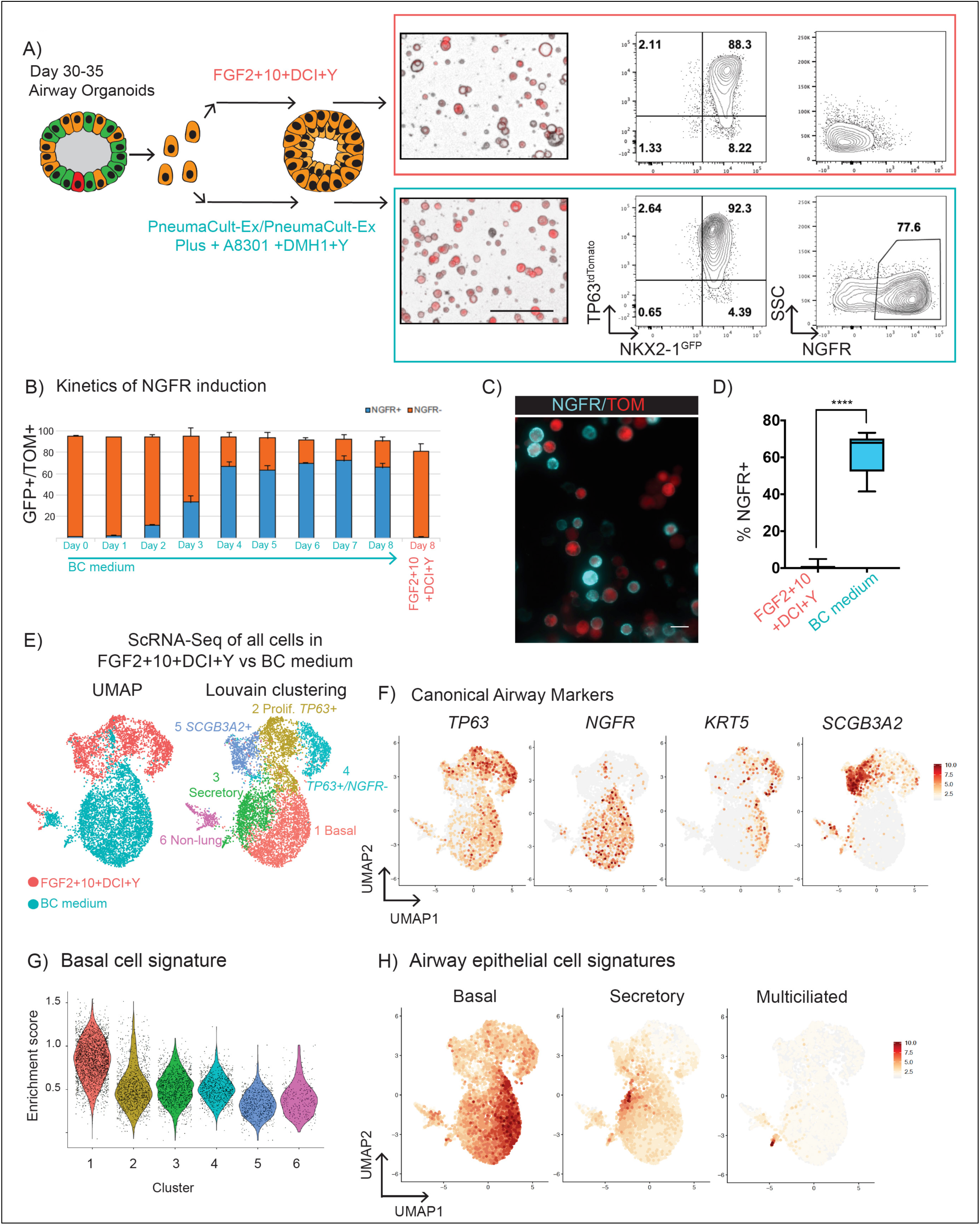
NKX2-1^GFP+^/TP63^tdTomato+^ cells adopt a molecular signature similar to primary basal cells. (A) Schematic of experiment: GFP+/TOM+ sorted cells were suspended in 3D Matrigel between days 30-35 of differentiation in FGF2+FGF10+DCI+Y or primary BC media. After 12-14 days morphology and tdTomato fluorescence were assessed (left panel) and the expression of NKX2-1^GFP^, TP63^tdTomato^ and NGFR quantified by flow cytometry (middle and right panel, representative plots). See supplemental figure 2 for additional details. (B) Kinetics of NGFR induction, quantified by flow cytometry in response to primary BC medium compared to continued FGF2+FGF10+DCI+Y over 8 days from day 33 to day 41 (n=3). (C) NGFR protein immunostaining (cyan) of BU3 NGPT cells in primary BC medium compared to TP63^tdTomato^ fluorescence (scale bar=20μm). (D) Percentage of NGFR+ cells quantified by flow cytometry between day 40-50 (n=12). (E) ScRNA-Seq of day 46 cells from primary BC or FGF2+FGF10+DCI+Y media. UMAP (left panel) displays the distribution based on culture conditions used to treat cells. Louvain clustering (res = 0.25) identifies 6 clusters (1-6). (F) UMAP of canonical BC (*TP63, NGFR, KRT5*) and SC (*SCGB3A2*) markers. (G) Graph of basal cell gene signature enrichment score for each cluster. (H) UMAP of the expression of basal, secretory and multiciliated cell gene signatures. See supplemental figures 3 and 4 for additional information.

Next, to determine whether the increase in NGFR expression in response to BC medium was coincident with a broader augmentation of a mature basal cell genetic program, we performed single-cell RNA-sequencing (scRNA-Seq). We differentiated BU3 NGPT iPSCs in our new protocol (Figure 2A), purifying GFP+/TOM+ cells on day 30-32 followed by replating for 3D culture in either “BC medium” vs continuing in FGF2+10+DCI+Y medium until scRNA-Seq analysis on day 46. Louvain clustering was applied to these combined samples and identified six populations (1-6, in descending order of size) (Figure 2E). Clusters 2,4 and 5 were predominantly composed of cells from FGF2+10+DCI+Y conditions whereas clusters 1 and 3 were almost entirely composed of cells from BC medium (Figure 2E). Based on the expression of canonical markers of lung and non-lung endoderm, cell-cycle and differentially expressed genes (DEG) in each cluster we annotated these clusters as follows; 1: “Basal”, 2: “Proliferative *TP63+*”, 3: “Secretory”, 4: “*TP63+/NGFR-*”, 5: “*SCGB3A2+*” 6: “Non-lung endoderm” (Figure 2E-F, S3A-E). Basal-like cells in both conditions (clusters 1,2 and 4) expressed markers including *TP63* and *tdTomato*, thus supporting the reporter specificity, and expressed key BC markers including *KRT13, KRT15, KRT17, AQP3, CAV1* and *FBLN1* (within the top 30 DEG in clusters 1,2 and 4 compared to 3,5 and 6) (Figure S3A-E). As expected *NGFR* was only expressed in the basal-like population cultured in BC medium (cluster 1; Figures 2F). Notably, we have previously demonstrated that plasticity of iPSC cells undergoing lung directed differentiation can sometimes result in reversion to non-lung endodermal lineages (McCauley et al. 2018; Hurley et al. 2019), however the day 30-32 sorting step for GFP+/TOM+ cells significantly reduced the percentage of non-lung endoderm cells in both conditions to less than 0.05% of all cells (Figure S3B).

For an unbiased assessment to test the correlation of these cells with genetic programs of airway cell types we generated benchmark gene signatures for primary human basal cells and their differentiated progeny. scRNA-Seq of freshly isolated primary adult human basal cells before (“P0”; after minimal culture manipulation) versus after differentiation in ALI cultures (4 days of submerged culture followed by 20 days of ALI) was performed according to established protocols (Figure S4) (Fulcher & Randell 2013). Similar to recent scRNA-Seq profiling of human tracheal epithelial cells, we identified populations of basal, secretory, and multiciliated cells and a rare population of ionocytes (Figure S4B-D) (Plasschaert et al. 2018). We generated gene-signatures composed of the top 30 DEGs, ranked by fold change for each of these cell types (Table 1) and applied the BC signature of primary cells to the iPSC-derived samples and determined cluster 1 (BC medium) had the highest BC enrichment score (“Basal” cluster =0.84±0.24, “Proliferative *TP63+*”=0.52±0.24, “Secretory”= 0.51±0.18 “*TP63+/NGFR-*” =0.51±0.15, and “*SCGB3A2+*”=0.32±0.15, mean enrichment score±SD) (Figure 2G-H). DEGs in the basal cluster included *KRT15, S100A2, CAV1* and *COL17A1*(Figure S3C) which were also highly expressed in primary BCs(Rock et al. 2009; Plasschaert et al. 2018). Taken together, our results indicate that when cultured in BC medium, GFP+/TOM+ cells proliferated and adopted a more mature airway BC program, including expression of NGFR accompanied by increased expression levels of a diversity of canonical basal cell markers. In view of their more basal-like transcriptional profile we designated GFP+/TOM+ cells in BC medium as iPSC-derived basal cells (“iBCs”). Of note, cluster 3 had the highest mean enrichment scores for secretory cell (SC) signature and a small subpopulation of cells within cluster 3 had multiciliated cell (MCC) gene signature. Canonical SC (*SCGB1A1*) and MCC (*FOXJ1*) markers were expressed in cluster 3 suggesting some differentiation was occurring in spheroids in BC medium (Figures 2H and S3B).

### iBCs Have Similar Stem Cell Properties to Primary Basal Cells

We next asked to what extent iBCs share the key stem cell properties of primary BCs: long-term self-renewal and multi-lineage differentiation (Figure 3A-G). In BC medium, iBCs were propagated for up to 10 passages or 170 days in 3D culture, while retaining GFP, TOM, and NGFR expression in the majority of cells (Figure 3B) and maintaining a normal karyotype (Figure S5A-B). One input iBC plated on day 45 yielded 6.9±1.7 × 10^6^ cells (mean±SD) 78 days later with a doubling time of 1.1 days (Figure 3B). To assess the differentiation and self-renewal of individual cells we adapted the 3D tracheosphere assay (Rock et al. 2009) which has been used to measure these properties for primary airway basal cells. We established a seeding density at which >95% of spheres were clonally derived from a single iBC (Figure S5C). iBCs were generated according to the basal cell protocol, NGFR+ cells were sorted, and plated in 3D culture at this clonal density in BC or differentiation “ALI” media. In both conditions, spheres formed and were composed of a stratified-appearing layer of predominantly NKX2-1+ cells (Figure 3C). In BC medium, NKX2-1+/TP63+ cells were readily identified in the outermost cell layer of the sphere. In differentiation medium, BCs, SCs and MCCs were identified in single spheres suggesting self-renewal and clonal multi-lineage differentiation of iBCs (Figure 3C). In the second approach we generated 2D ALI cultures from iBCs. On day 46, NGFR+ iBCs were sorted and plated on Transwell inserts in differentiation “ALI” medium and compared to presorted cells (Schematic Figure 3A). Without NGFR sorting, we observed patchy differentiation in Transwell cultures, evident as frequent areas devoid of staining for the above lung markers and some areas devoid of TOM expression (Figure 3D). In contrast, NGFR+ sorted cells initially formed a confluent TOM+ epithelial layer in submerged culture and after 16 days in air-phase culture SCs (SCGB1A1+, MUC5AC+) and beating MCCs were distributed homogenously across the entire Transwell (Figure 3E and Supplemental Video 1). Confocal microscopy and transverse sections confirmed that NGFR+ sorted cells had formed pseudostratified epithelia composed of MCCs (ACT+, FOXJ1+) and SCs (SCGB1A1+, MUC5AC+) (Figure 3E-G). Importantly, after ALI differentiation, iBCs had also formed TP63+/KRT5+/NGFR+ cells with the expected “basal” location of bona fide BCs, along the basal membrane (Figure 3G). To assess integrity of this epithelium, we measured TEER. Both NGFR sorted and GFP+/TOM+ cells formed epithelial layers with similar TEER to primary controls and significantly higher TEER than ALI differentiations prepared in parallel from GFP+/TOM+ cells that had been maintained in FGF2+10+DCI+Y medium without BC medium maturation (Figure 3H). We also tested the extent to which the multi-lineage differentiation capacity of iBCs was maintained after extended *in vitro* expansion in BC medium. After 10 passages of expansion, iBCs retained their capacity to form trilineage, pseudostratified airway epithelium when transferred to ALI culture (Figures S5D). After cryopreservation and subsequent thaw iBCs retained their expression of basal cell markers, proliferated in BC medium and retained trilineage differentiation potential in ALI conditions (Figure 3I). Taken together these results indicated successful derivation from iPSCs of iBCs which exhibit the airway stem cell properties of self-renewal and multipotent differentiation.

**Figure 3:**
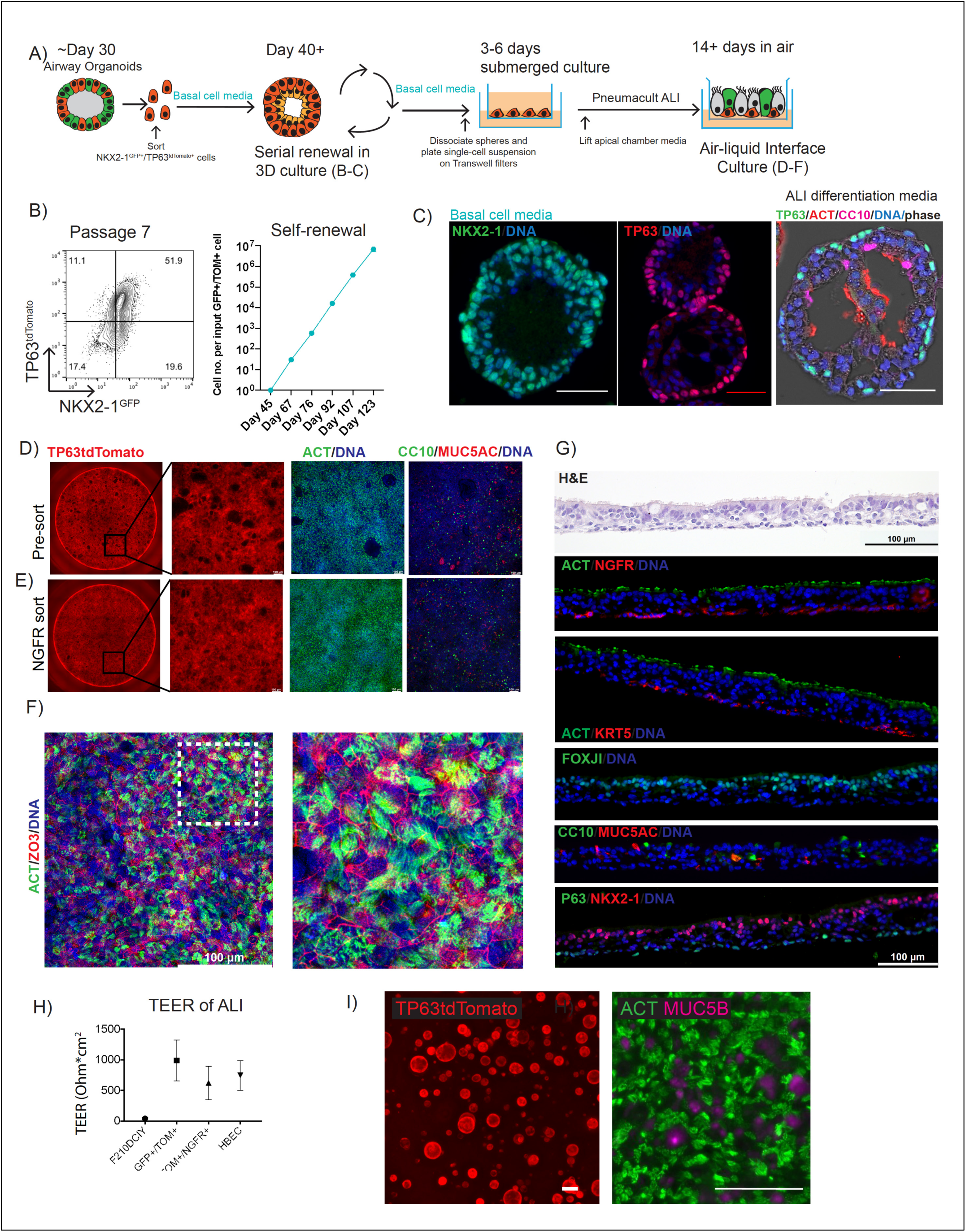
iBCs undergo self-renewal and multi-lineage differentiation. (A) Schematic of experiment: GFP+/TOM+ cells sorted on approximately day 40 of differentiation expanded in BC medium and then characterized in terms of self-renewal and multi-lineage differentiation in air liquid interface (ALI). B) Representative flow cytometry plots of NKX2-1^GFP^ vs TP63^tdTomato^ expression in cells at passage 7, following expansion in 3D culture with BC medium (left panel). Cell numbers per input GFP+/Tom+ cell up to 123 days of directed differentiation (right panel). (C) Immunolabeling of representative spheroid on day 83 of differentiation in either BC medium (left and middle panel) or after 10 days in ALI differentiation medium (right panel) with antibodies against indicated markers (nuclei stained with Hoechst; scale bar=50μm). See also supplemental figure 5C. (D-E) GFP+/TOM+ cells were expanded in BC medium until day 46 and plated on Transwells with (E) or without (D) GFP/TOM/NGFR sorting. Representative images of the endogenous TP63^tdTomato^ fluorescence during submerged culture of BU3 NGPT-derived ALI. Stitched image of whole Transwell insert (Ø =6.5mm) (1^st^ column) and zoom-in (2^nd^ column). Immunolabeling antibodies against ACT, CC10, and MUC5AC after 16 days of ALI culture. (3^rd^ and 4^th^ columns) (DNA stained with DRAQ5; sScale bar =100μm). (F) Confocal microscopy of BU3 NGPT-derived ALI cultures immunolabeled with antibodies against ACT and ZO3 (DNA stained with DRAQ5; scale bar =100μm). (G) Transverse section of BU3 NGPT-derived ALI cultures shown in E and stained with hematoxylin and eosin or antibodies against ACT, NGFR, KRT5, FOXJ1, CC10, MUC5AC, P63, and NKX2-1 (DNA stained with DAPI; scale bar=100μm). (H) TEER measurements of Transwell ALI cultures, comparing GFP+/TOM+ from FGF2+10+DCI+Y medium (n=5), BC medium with (N=4) or without (n=5) NGFR sorting and compared to primary HBEC controls (n=21). (I) TP63^tdTomato^ fluorescence in BU3 NGPT iBCs after cryopreservation and thaw (scale bar =200μm) (left panel). Immunolabeling of ALI cultures generated from cryopreserved iBCs with antibodies against MUC5B and ACT (scale bar=100μm).

### Single-cell RNA Sequencing of iBC-derived Airway Epithelium

To more completely assess the molecular phenotypes of the differentiated cell types generated from iBCs (Figure 4A), we performed scRNA-sequencing of ALI cultures from NGFR sorted cells. Louvain clustering identified five clusters (1-5) with gene expression profiles reminiscent of airway epithelial cell types and one proliferative cluster (6) (Figure 4A-E, S6A-B). We found no discrete cluster of non-lung endoderm cells (low to undetectable *CDX2, AFP, ALB*). We compared the top 20 DEGs in each cluster to the single-cell RNA sequencing dataset of differential gene expression in primary airway epithelial cell types (Figure 4B, S4 and Table 1) and published datasets (Plasschaert et al. 2018) and annotated the cell types as follows: “Secretory” (cluster 2, DEGs: *SCGB1A1, SCGB3A2, MUC5B, C3, XBP1, CEACAM6, CXCL1*), “Basal” (cluster 6, DEGs: *KRT5, KRT17, S100A2*), “Intermediate” (cluster 1, DEGs: *KRT4, KRT12,KRT13*, and *KRT*15), “Immature MCC” (cluster 3, DEGs: *CCNO, CDC20b, NEK2*) and “MCC” (cluster 4, DEGs: *TUBB4B, TUB1A1, TPP3* and *CAPS*) (Figures 4A-E, S6A-B). Enrichment scores for gene-signatures of primary BCs, SCs and MCCs correlated with these annotated clusters with enrichment scores of 0.77±0.16, 0.54±0.26 and 1.54±0.27 (mean enrichment score±SD), respectively (Figures 4D, S6A). FOXJ1, a key transcription factor expressed in MCCs, was found enriched in both ciliated cell clusters (Figure 4C); however, FOXN4, recently identified as a marker of immature primary human airway MCCs, was found only in the immature iPSC-derived MCCs (Figure S6A) (Plasschaert et al. 2018). The expression levels of key airway epithelial markers in iBC-derived ALI cultures was confirmed using qRT-PCR and compared to primary controls (Figure 4F). There was no evidence to suggest the presence of ionocytes, neuroendocrine or alveolar epithelial cells amongst the 3,500 iPSC-derived cells analyzed (data not shown). In summary, pseudostratified airway epithelium derived from iBCs is composed of BCs, SCs and MCCs.

**Figure 4:**
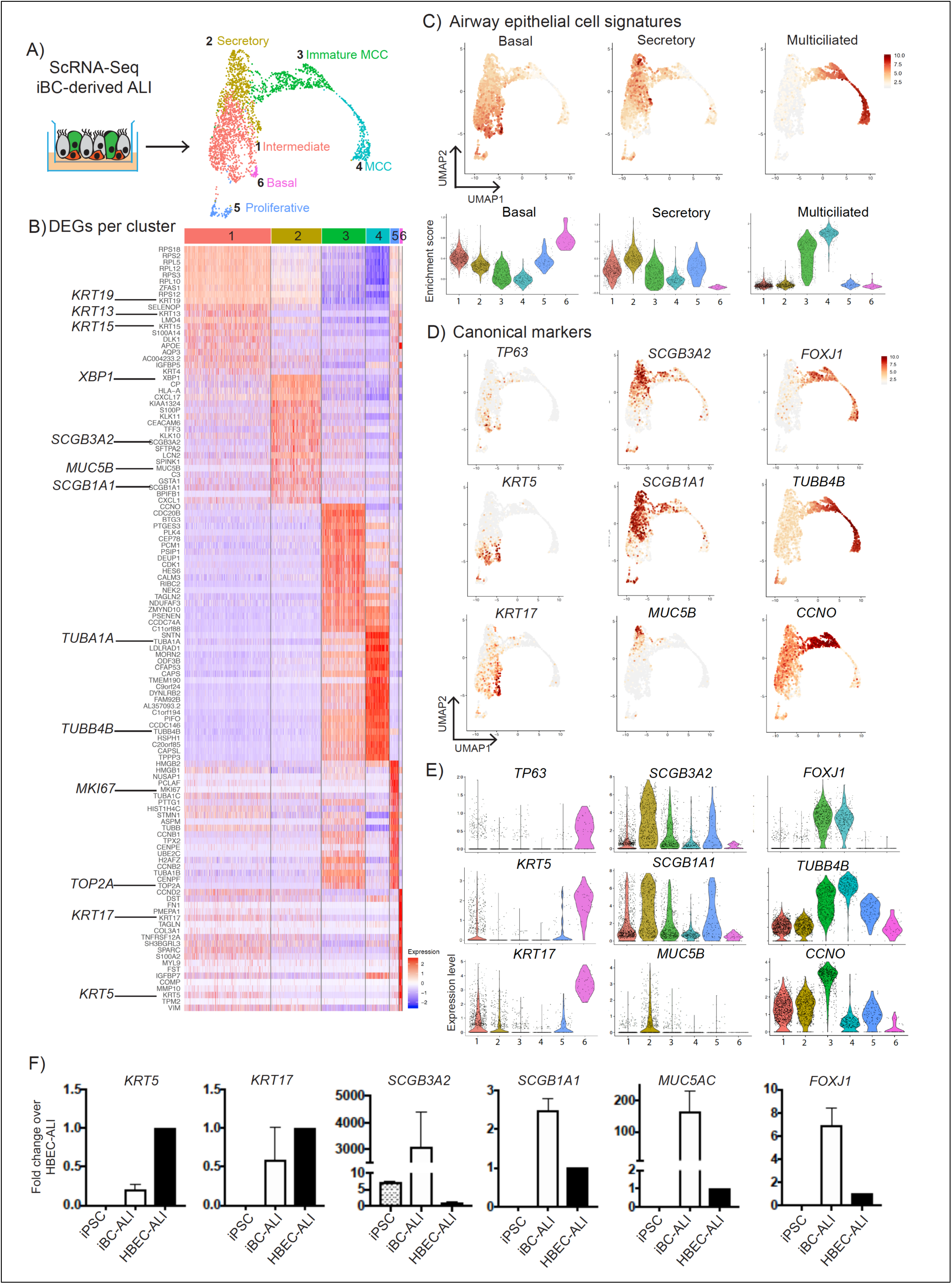
scRNA-Seq profiling of iBCs and their differentiated progeny. (A) Schematic of scRNA-Seq experiment and UMAP with Louvain clustering (clusters 1-6) of iBC-derived ALI. See also supplemental figures 4 and 6. (B) Top 20 differentially expressed genes (DEG) per cluster. (C) UMAPs of primary BC, SC and MCC gene-signatures applied to iBC-derived ALI (upper row). Violin plots of the enrichment score of clusters 1-6 for BC, SC and MCC gene signatures (lower panel). (D) UMAPs of the canonical BC (*TP63, KRT5, and KRT17*), SC (*SCGB3A2, SCGB1A1, and MUC5B*) and MCC (*FOXJ1, TUBB4B, and CCNO*) across clusters 1-6. (E) Violin gene-expression plots of same panel of markers shown in (D). (F) qRT-PCR validation of key airway markers in iBC-derived ALI (iBC-ALI) compared to undifferentiated iPSCs and primary HBEC-derived ALI (HBEC-ALI). Fold change is 2^-ΔΔCt^ normalized to HBEC-ALI.

### Function and Physiology of iBC-derived Airway Epithelium

Human airway epithelium has diverse biologic and physiologic roles, some of which are recapitulated in primary human airway cultures. Regulation of ion-flux by CFTR and mucus cell metaplasia in response to the inflammatory cytokine interleukin 13 (IL-13) are examples. Hence, we sought to determine the extent to which iBCs and their differentiated progeny reproduced these features. First, CFTR-dependent current, representing ion-flux regulated by apically-localized CFTR, was measured using the gold-standard Ussing chamber approach. First we blocked ENaC channels with amiloride followed by CFTR stimulation with forskolin (Fsk) and VX-770, a CFTR potentiator, with subsequent CFTR inhibition using CFTRinh-172 (Figure 5A). In ALI from Figure 3E, Fsk-activated short-circuit currents (ΔIsc) were 30.5 ± 3.4 μA/cm^2^ and VX-770-activated ΔIsc were 0.4 ± 2.0 μA/cm^2^, which were then inhibited by the CFTR specific inhibitor (ΔIsc = 30.1 ± 2.2 μA/cm^2^) (Figure 5B). These values were in a similar range to published data using primary human airway cultures (Gentzsch et al. 2016). Second, we tested whether iBC-derived ALIs could recapitulate the mucus cell metaplasia seen in asthma. IL-13 is an inflammatory cytokine in asthma and can induce mucus metaplasia in primary human airway epithelial cell cultures (Seibold 2018). In response to IL-13 added on day 10 of ALI differentiation of iBCs, we found a significant 1.75 fold increase in numbers of MUC5AC+ cells (p=0.002), increased expression of the goblet cell transcriptional regulator *SPDEF*, and a decrease in *MUC5B* expression (Figures 5C-E). These findings are in keeping with primary mouse and human adult airway epithelial cultures treated with IL-13 where a number of studies have found increased MUC5AC+ cell numbers, via the activation of STAT6 and SPDEF, at the expense of MCCs and MUC5B cell frequencies (Seibold 2018; Kondo et al. 2002; Woodruff et al. 2009). Together these data suggest that iPSC-derived airway epithelium shares key physiologic and biologic features with human airway epithelium.

**Figure 5:**
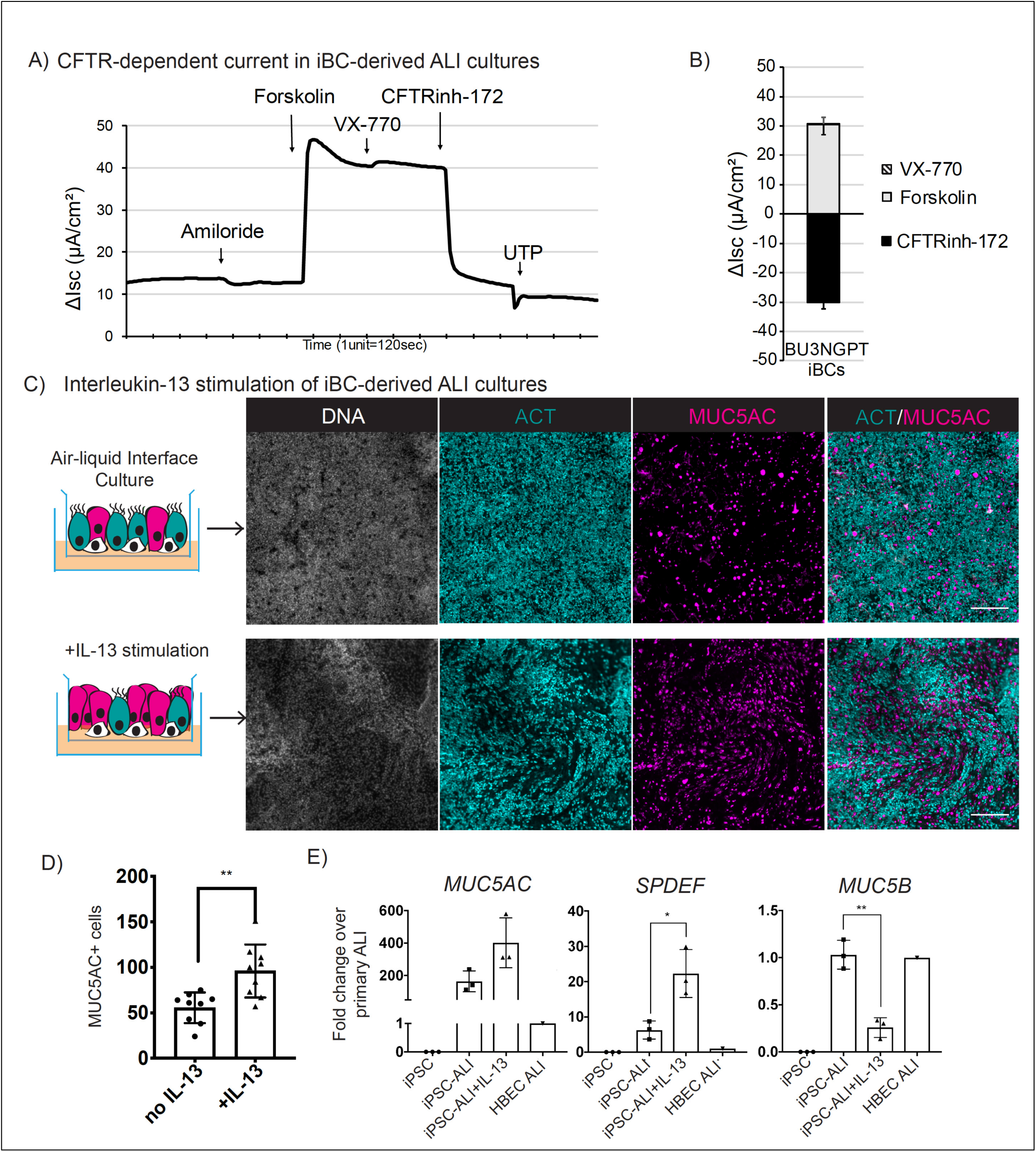
iBC-derived airway epithelium recapitulates key physiologic and functional properties of human airways. (A) Representative electrophysiological traces from Ussing chamber analysis of BU3 NGPT-derived ALI cultures shown in Figure 3E. (B) Mean and SD of electrophysiological values from (A) (n=3). (C) Representative images of BU3 NGPT iBC-derived ALI cultures, with or without IL-13 treatment, immunolabeled with antibodies against ACT and MUC5AC (DNA stained with Hoechst; scale bar=200μm). (D) Quantification of the number of MUC5AC+ cells per high power field for IL13 treated vs untreated wells (n=3). (E) qRT-PCR quantification of mRNA expression levels of *MUC5AC, SPEDF* and *MUC5B* in IL-13 treated vs untreated cells compared to primary untreated ALI controls (n=3). (*=p<0.05, **=p<0.01).

### iBCs Generated from Patient-Specific iPSCs Model Cystic Fibrosis and Primary Ciliary Dyskinesia

To expand the application of iBCs for disease modeling we first sought to adapt our approach so that a diversity of normal or patient-specific iPSC lines could be differentiated into iBCs without requiring the use of knock-in fluorochrome reporters. We found through repeating our protocol with the BU3 NGPT line, we could replace GFP+/TOM+ sorting with antibody-based sorting, initially gating on NGFR+ cells to prospectively isolate putative iBCs (Figure 6). For example, cells grown until days 40-42 with our protocol (Figure 6A-C), were 73.7±5.9% (mean±SD) GFP/TOM double positive, and gating on NGFR+/EpCAM+ cells yielded a population of cells highly enriched in GFP/TOM co-expressing cells (94.3±1.1%,mean±SD) (Figure 6B). A representative experiment is shown in Figure 6B; 73.6% of cells were GFP+/TOM+, 55.3% NGFR+/EPCAM+, and 94.5% of sorted NGFR+/EPCAM+ cells were GFP+/TOM+ (Figure 6B) Sorting solely on NGFR+ or NGFR+/EpCAM+ and replating cells in ALI cultures resulted in successful derivation of a TOM+ retaining, pseudostratified airway epithelium, similar to sorting GFP+/TOM+ cells and in contrast to unsorted controls (Figure 6C). Four additional iPSC lines (DD001m, PCD1, 1566, 1567) and an ESC line (RUES2) were also differentiated, recapitulating our basal cell differentiation protocol while completely replacing the need for fluorescent reporters through the use of CD47^hi^/CD26^neg^ sorting on day 15 to purify NKX2-1+ progenitors as we have published(Hawkins et al. 2017), followed by NGFR+ sorting after day 40 to purify candidate iBCs (see methods and schematic Figure 6A). NGFR+ cells from all five iPSC/ESC lines differentiated into pseudostratified epithelia composed of MCCs, SCs and BCs (Figure 6D-G).

**Figure 6:**
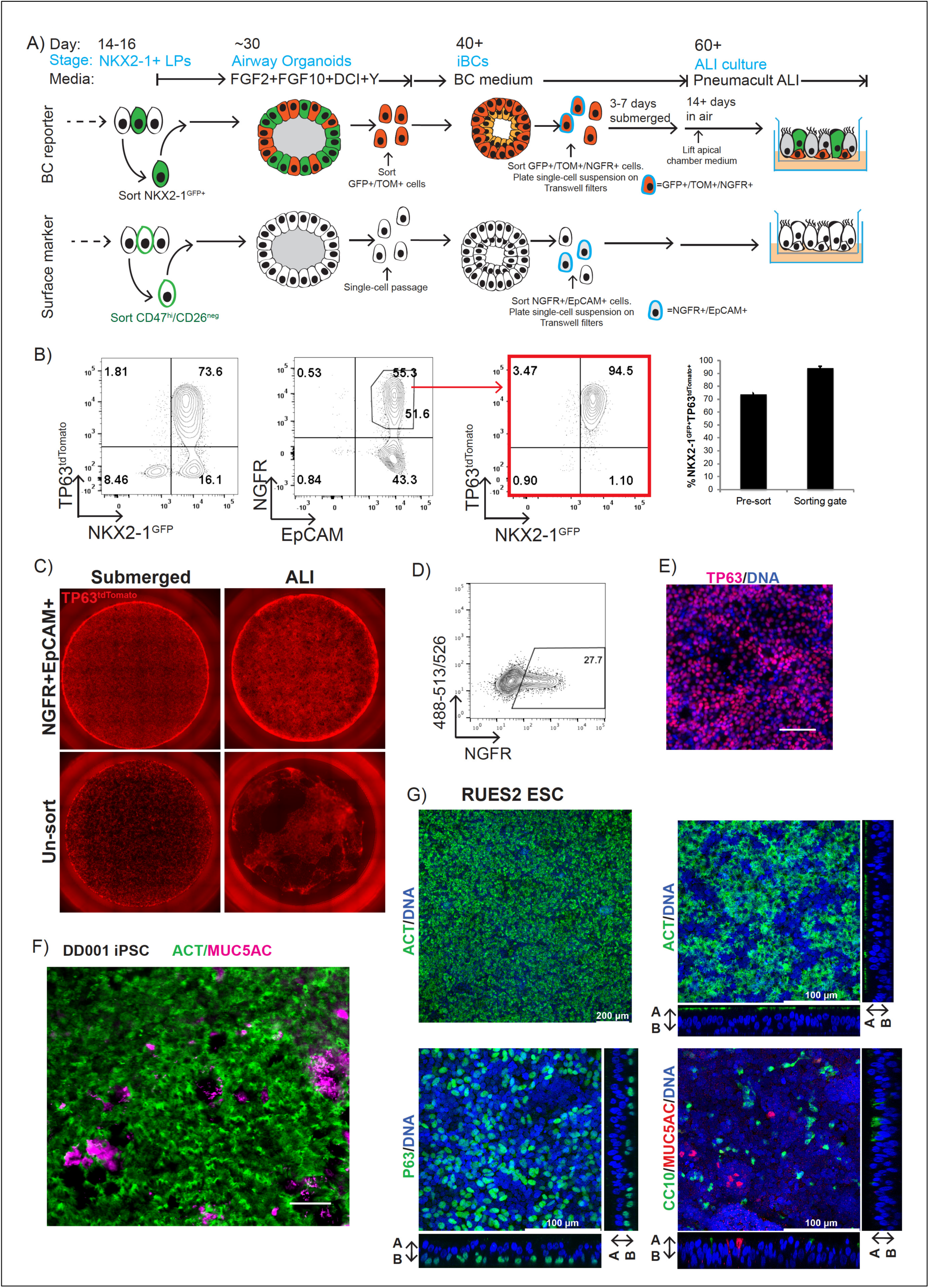
A surface marker strategy for purifying iBCs using NGFR replaces the need for fluorescent reporters. (A) Schematic of NKX2-1^GFP^/TP63^tdTomato^ reporter vs surface marker iBC protocols. (B) Representative flow cytometry plots of BU3 NGPT iBCs on day 40 of differentiation and labeled with antibodies against NGFR and EpCAM. Red arrow indicates sorted NGFR+/EpCAM+ cells are 94.5% GFP+/TOM+. Enrichment is quantified in the right panel (n=3). (C) Representative images of the endogenous TP63^tdTomato^ fluorescence in whole Transwell filters (Ø =6.5mm) seeded with sorted (NGFR+/EpCAM+) or unsorted cells after one week in submerged culture followed by lifting medium from the apical chamber for two weeks (ALI). (D) Representative flow cytometry plot of a non-reporter iPSC line stained for NGFR is shown. (E) Representative image of sorted NGFR+ cells plated on Transwell filters and after 7 days immunolabeled with an anti-TP63 antibody (lower right panel; nuclei stained with XX; scale bar=100μm). (F) Confocal microscopy of DD001m iPSC-derived ALI cultures immunolabeled with antibodies against ACT and MUC5AC (scale bar=200μm). (G) Representative image of RUES2 ESC-derived ALI immunolabeled with an anti-ACT antibody (nuclei stained with DRAQ5; scale bar=200μm) (Upper left) and confocal microscopy of RUES2 ESC-derived ALI cultures immunolabeled with antibodies against ACT, TP63, CC10 and MUC5AC (nuclei stained with DRAQ5; scale bar=100μm).

Next we applied our antibody-based iBC purification protocol for modeling two genetic airway diseases, cystic fibrosis (CF) and primary ciliary dyskinesia (PCD) (Figure 7A). We identified a CF patient homozygous for the most common *CFTR* mutation (c.1521_1523delCTT, p.Phe508del) and a PCD patient homozygous for a mutation in *DNAH5* (c.12617G>A, p.Trp4206Ter), one of the most common genes in PCD, and generated iPSCs from these individuals by reprogramming peripheral blood specimens (see methods and Figure S7A-B). For cystic fibrosis modeling we first used CRISPR based gene-editing to correct F508del *CFTR* mutation in both alleles and then differentiated pre- and post-gene corrected paired syngeneic iPSC clones (hereafter CF and Corr CF) in our protocol using our antibody-only based sorting methods. We derived iBCs from each line, performed NGFR sorting (Figure 6A) and differentiated the sorted cells in ALI cultures, producing airway epithelia composed of BCs, SCs and MCCs (Figure S7C). As expected Ussing chamber analysis indicated minimal CFTR-dependent current in CF ALI cultures (Forskolin ΔIsc =0.8μA/cm^2^, CFTRInh-172 ΔIsc =-0.8μA/cm^2^). In marked contrast, correction of the F508del mutation in one allele led to restoration of CFTR-dependent current similar in magnitude to BU3 NGPT (non-CF) or published data of low-passage non-CF primary airway epithelial cells (Forskolin ΔIsc =35.1±1.77μA/cm^2^, CFTRInh-172 ΔIsc =43.2- ±2.5μA/cm^2^, Figure 7B-C) (Gentzsch et al. 2016).

**Figure 7:**
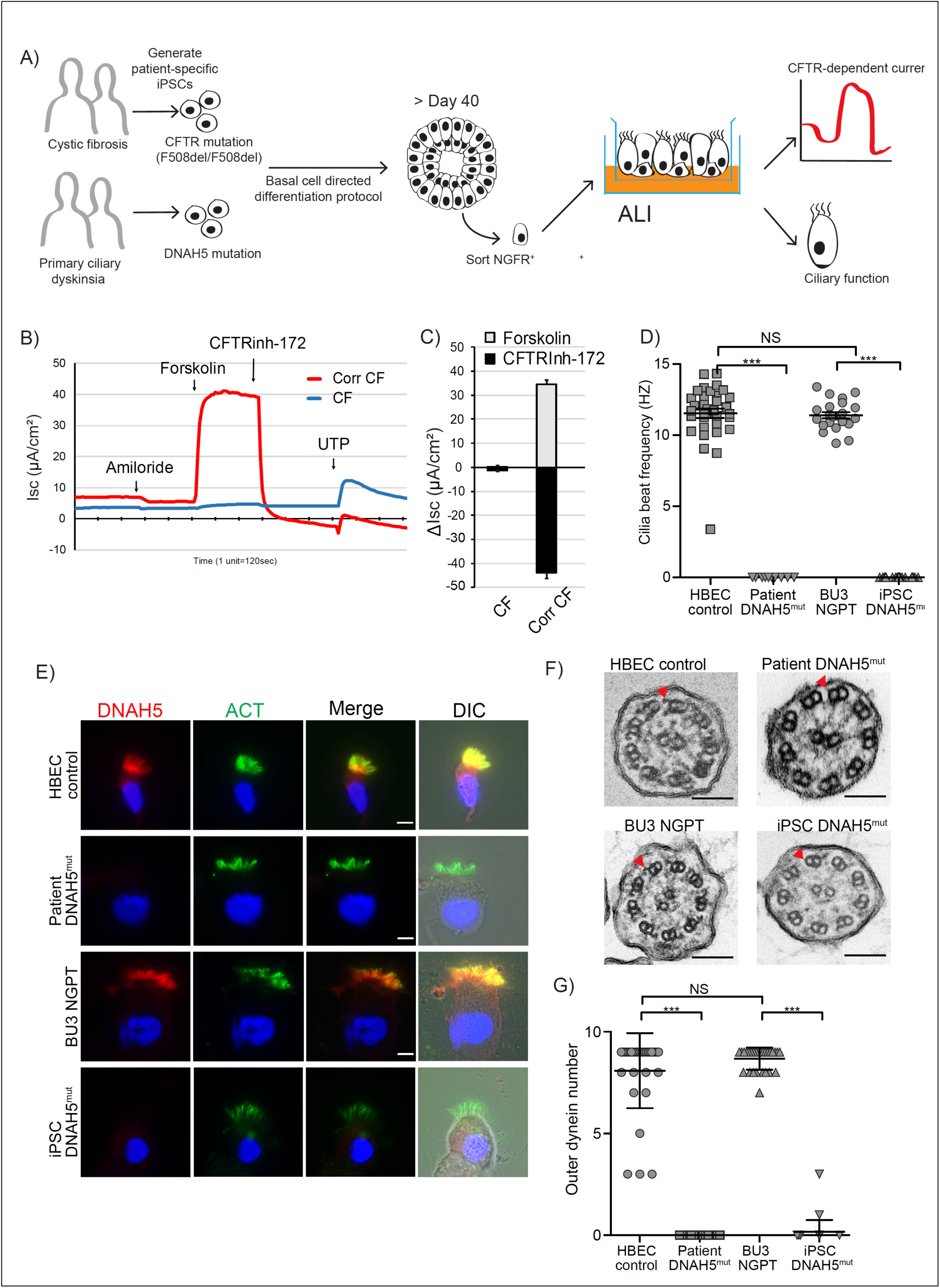
iBCs enable *in vitro* modeling of cystic fibrosis and primary ciliary dyskinesia. (A) Schematic of CF and PCD disease modeling experiments, relying on surface markers. See also supplemental figure 7. (B) Representative electrophysiological traces from Ussing chamber analysis of ALI cultures generated from CF iPSCs and corrected CF iPSCs (Corr CF). (C) Mean and SD of electrophysiological values from (B) (n=3). (D) Ciliary beat frequency in ALI cultures generated from *DNAH5* mutant nasal epithelial cells, *DNAH5* mutant iPSCs, primary HBECs and wild-type iPSCs (BU3 NGPT). (***=p<0.001). (E) Immunolabeling of multiciliated cells from the samples detailed in (D) with antibodies against ACT and DNAH5 (nuclei stained with DAPI; scale bar=10μm). (F) Transmission electron microscopy of cilia from samples detailed in (D). (G) Quantification of the number of outer dynein arms detected in cross sections of cilia from samples detailed in (D). (***=p<0.001)

To develop a model for primary ciliary dyskinesia, we differentiated our PCD iPSC line carrying the *DNAH5* mutation in our 3D culture protocol and purified putative iBCs by NGFR sorting. After replating the sorted cells in ALI cultures to generate MCCs, we compared these cells to primary cell controls consisting of differentiated MCCs prepared from the primary nasal epithelium of the PCD donor. We included additional MCC controls generated from BU3 NGPT iBCs or from normal primary HBECs. By visual inspection using phase microscopy, abundant cilia beating was observed in BU3 NGPT cultured cells, however, none was observed in *DNAH5* mutant iPSC cells or in cultured primary nasal cells obtained from the PCD subject. To quantify this difference, we measured ciliary beat frequency (CBF) in each culture. Cilia motility was not detected in the MCCs generated from either the *DNAH5* mutant iPSC line or from *DNAH5* mutant primary cells, compared to normal controls (Figure 7D and supplemental videos 2A-D). The possibility that *DNAH5* mutant iPSCs had failed to produce cilia was excluded by immunofluorescent staining with antibodies against ACT, which confirmed the widespread presence of normal appearing cilia in all samples. In addition, increased tethering of MUC5AC+ mucus strands was observed on the apical surface of the differentiated epithelium generated from the mutant iPSCs (Figure S7D).

To characterize the ciliary defect in *DNAH*5 mutant iPSC, we performed immunofluorescent staining with an antibody against DNAH5 protein. *DNAH5* encodes the heavy chain 5 dynein motor protein found in the multiprotein complex that composes the outer dynein arm (ODA) of the ciliary axoneme. Mutations in *DNAH5* typically render the cilia severely hypokinetic due to loss of the entire ODA structure. In MCCs generated from normal iPSCs and normal HBECs, DNAH5 protein was present and colocalized with ACT along the length of cilia (Figure 7E). In MCCs cells generated from *DNAH5* mutant cells, DNAH5 protein was not detected in either iPSC-derived or primary nasal-derived MCCs cells. In contrast DNALI1, an inner dynein arm (IDA) protein, was detectable by immunostaining, suggesting an intact IDA complex (Figure S7E). Transmission electron microscopy of *DNAH5* mutant iPSC-derived MCCs cells showed lack of the ODA, identical to the defect in cilia of nasal cells obtained from the patient, and in contrast to MCCs cells obtained from control iPSC and normal HBEC (Figure 7F-G). Taken together these results indicate that patient-specific iBCs purified without using fluorescent protein reporters can be successfully applied to model a variety of airway epithelial diseases, including CF and PCD.

## Discussion

Here we report the differentiation of iPSCs into cells that are molecularly and functionally similar to the main stem cell of human airways, the basal cell. The derivation of a tissue resident stem cell from human iPSCs has considerable implications for regenerative medicine research as it overcomes several important hurdles currently limiting progress in this field.

For example, directed differentiation protocols are often lengthy, complex, and yield immature and heterogeneous cells. Frequently these issues are due to lack of precise knowledge regarding developmental roadmaps associated with cellular embryonic origin. As a result, the goal of deriving specialized human cell types that are molecularly and functionally similar to their endogenous counterparts has been hindered. Despite these issues, significant progress has been made in recent years in deriving lung epithelial cells from iPSCs (Huang et al. 2014; Firth et al. 2014; Gotoh et al. 2014; Dye et al. 2015; Miller et al. 2018; Hawkins et al. 2017; McCauley et al. 2017; Jacob et al. 2019). The identification of surface markers or development of fluorescent reporter iPSC lines can overcome the diversity of cell types generated during directed differentiation. These approaches in combination with organoid-based cultures have resulted in the derivation of several lung epithelial cell types and a number of groups, including ours, have generated populations of lung cells similar to secretory, multiciliated, neuroendocrine, and alveolar type 2 cells (Hawkins et al. 2017; Gotoh et al. 2014; Konishi et al. 2016; Jacob et al. 2019; Miller et al. 2018; Chen et al. 2017). TP63+ populations derived from iPSCs previously have been observed stochastically and their characterization was based only on the expression of a handful of canonical markers limiting any conclusions as to whether basal-like cells had been produced (Hawkins et al. 2017; McCauley et al. 2017; Chen et al. 2017; Konishi et al. 2016; Dye et al. 2015). Without the ability to purify these cells from the heterogeneous mix of other iPSC-derived lineages, little progress has been made phenotyping or testing their functional potential. Here we report several advances that culminate in the efficient derivation and purification of basal cells from iPSCs. Using a dual fluorescence basal cell reporter iPSC line, we recapitulated *in vitro* the milestones of *in vivo* airway development and basal cell specification (Y. Yang et al. 2018). First, during the emergence of the earliest detectable lung epithelial program, rare TP63+ cells were detected. Next, in response to the withdrawal of Wnt signaling and in the presence of FGF2 and FGF10, an immature airway program with upregulation of TP63 and SCGB3A2 was evident. In response to primary basal cell medium and inhibition of SMAD signaling, these TP63+ cells matured and adopted molecular and functional phenotypes similar to basal cells, including the capacity for extensive self-renewal for over 150 days in culture and trilineage airway epithelial differentiation. We conclude that the resulting cells fulfill the criteria to be termed iPSC-derived basal cells or iBCs.

From a practical perspective, insight and methodologies developed herein should expand the application of iPSC technology. The shared properties of primary BCs and iBCs for expansion and differentiation in ALI culture makes them attractive for a variety of *in vitro* or *in vivo* applications. Specifically, the ability to purify iBCs based on the surface marker NGFR, expand the resulting cells in 3D culture for at least 10 passages and cryopreserve cells should minimize variability associated with directed differentiation and can potentially offset the cost and time required for these protocols.

In terms of the utility of this platform for disease modeling, we selected two genetic diseases with unique research challenges. In CF, where many mutations in one gene necessitate individual models of disease for predicting personalized therapeutics, we demonstrated that patient-specific iPSCs or their gene-edited progeny can be differentiated into iBCs and give rise to airway epithelia exhibiting quantifiable CFTR-depended currents of sufficient magnitude for disease modeling using the gold-standard Ussing chamber assay of CFTR function. One potential application of CF iBC-derived epithelium may be in generating a sufficient number of cells for high throughput screening of compounds for *CFTR* mutations including nonsense variants. A second application may be in predicting *in vivo* responses of patients to specific CF drugs. Finally, it is possible that autologous *CFTR*-corrected iPSC-derived iBCs may eventually be employed therapeutically for *in vivo* regeneration of airway tissue. For PCD, where many mutations in many genes controlling ciliogenesis can contribute to the disease, we determined that iBC-derived MCCs model both the functional and ultrastructural defects observed in *DNAH5* mutant primary donor derived cells. These proof-of-concept experiments suggest that the iPSC platform may be helpful in determining mechanisms of pathogenicity for genetic airway diseases and may serve as a platform to develop novel therapeutics.

While we conclude that iBCs share essential features with their endogenous counterparts, several questions remain including the maturity status, regionality and *in vivo* competence of iBCs. Typically, iPSC-derived specialized cell types are at fetal stages of development (Studer et al. 2015). A detailed characterization of the fetal and adult basal cell program is currently lacking for human airways. Hence, it is not yet possible to precisely benchmark the developmental stage of iBCs (Nikolić et al. 2018). Human airways change in appearance and cellular composition from proximal to distal; the trachea is pseudostratified and as airway caliber narrows, the epithelium eventually becomes simple columnar, suggesting regional patterning that is likely maintained by regional BCs. There is insufficient data using primary human airways to determine if regional patterning of BCs exists and to address whether iBCs are biased toward a particular region of the airway. We note differences in the pattern of expression of two secretoglobins, *SCGB3A1* and *SCGB3A2*, in iPSC-derived cells compared to primary cells. *SCGB3A1*, predominantly expressed in the larger airway, is the most highly expressed gene in primary BC-derived secretory cells while *SCGB3A2* is expressed at very low levels. iBC-derived secretory cells express high *SCGB3A2* and low/absent *SCGB3A1*. These differences further raise the question of immaturity and/or proximal/distal patterning (Reynolds et al. 2002). In terms of the specialized cell types derived from BCs there are additional unresolved questions. We did not observe rare airway epithelial cells, such as ionocytes or brush cells, in our ALI cultures. It is not yet known when or through which mechanism these cells emerge during development in human airways, and our iBC system does not yet provide a model for studying these rare cell types.

Lastly, there remains uncertainty as to whether iBCs have greater potential for stable expansion in culture than primary BCs. The differentiation capacity of primary BCs is known to deteriorate with serial passaging and with it their utility for disease modeling (Gentzsch et al. 2016). While iBCs still differentiate into airway epithelial cells after 10 passages, it is possible that, as with primary BCs, extended passage also has an effect on the transcriptome, epigenome and differentiation capacity of iBCs. Further studies will be required before any conclusions can be drawn regarding comparative stability in culture. That said, efficient derivation of an identifiable, tissue-specific stem cell, capable of long-term self-renewal, cryopreservation capacity, and retained multilineage differentiation overcomes many of the hurdles that face iPSC technology for studies of respiratory mucosa and diseases of pulmonary epithelium.

## Author contributions

FJH, SS, BRD and DNK conceived the work, designed the experiments, and wrote the manuscript. SS and BRD designed the targeting strategy and generated the TP63^tdTomato^ iPSCs. FJH, SS, BRD and DNK developed the basal cell differentiation protocol. FJH, SS, CB, AB, MLB, AMC, RW and JLS performed differentiation experiments. CV performed bioinformatic analysis of scRNA-sequencing. YT and TMS generated CF iPSCs and performed gene-correction. RW generated ALIs from CF iPSCs. AR and EJS performed Ussing chamber ananlysis of ALIs. SXH provided critical input. SHR provided primary HBECs and critical input. AH and SLB performed the experiments characterizing cilia in iPSC-derived MCCs including TEM and CBF.

## Acknowledgements

We thank Brian R. Tilton of the BUSM Flow Cytometry Core and Yuriy Alekseyev of the Boston University School of Medicine (BUSM) Single Cell Sequencing Core; both supported by NIH grant 1UL1TR001430. We also acknowledge Dr. Zhengmei Mao of UTHealth Microscopy Core Facility for histological sample preparation. For facilities management, we are indebted to Greg Miller (CReM Laboratory Manager) and Marianne James (CReM iPSC Core Manager) supported by grants R24HL123828 and U01TR001810. For primary HBECs we thank The Marsico Lung Institute Tissue Procurement and Cell Culture Core, University of North Carolina, Chapell Hill, NC. The current work was supported by R01HL139799 and Cystic Fibrosis Foundation (CFF) grant HAWKIN17I0 to FJH R01HL095993; Harry Shwachman Cystic Fibrosis Clinical Investigator Award, NIH CHRC K12HD052896 and Alfred and Gilda Slifka Fund to RW; TS was supported by grant U54 DK110805; American Thoracic Society Foundation, Primary Ciliary Dyskinesia Foundation, Kovler Family Foundation awards to (AH); CFF grants SORSCH 13XX0 and SORSCH 14XX0 to EJS; R01HL146601 and R01HL128370 to SLB; CFF DAVIS15XX1, DAVIS17XX0 to BRD; NIH R01 HL139876 to BRD and EJS; R01HL122442, R01HL095993, U01HL134745, U01HL134766 to DNK, U01HL148692 to DNK and FJH.

