## Supplemental methods, figures and tables for "Derivation of Airway Basal Stem Cells from Human Pluripotent Stem Cells"

### Supplemental Figures

Supplementary Figure 1: Characterization of BU3 NKX2-1$^{GFP}$;TP63$^{tdTomato}$ iPSC line

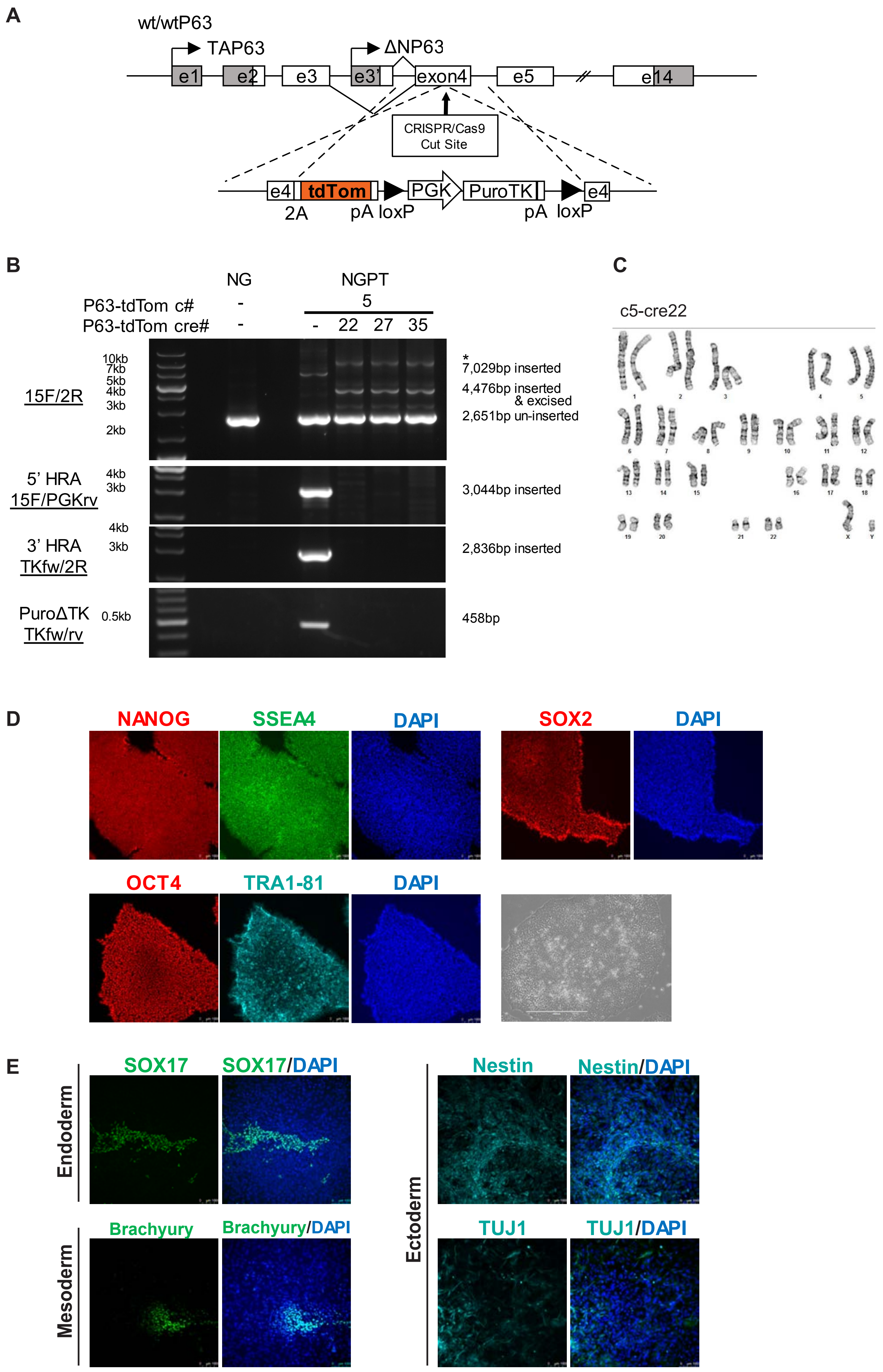

**Supplemental figure 1: Characterization of BU3 NKX2-1$^{GFP}$;TP63$^{tdTomato}$ iPSC line.**

A) Schematic of targeting strategy to introduce a 2A-tdTomato cassette at a double-stranded break site within exon 4 of one allele of *TP63*. The donor vector includes a floxed PGK promoter-driven antibiotic selection cassette (puroTK, consisting of a fused Puro resistance-thymidine kinase [TK] cassette) which is excised following transient Cre recombinase exposure. Negative selection was performed by adding ganciclovir to kill clones without successful PuroR-TK cassette excision.

B) *TP63* locus targeting screening by inside-outside PCR of gDNA from iPSCs

C) Karyotyping of BU3 iPSC line with successful monoallelic integration of the 2A-tdTomato cassette (clone 5) and after Cre-mediated excision of the antibiotic selection cassette (clone 22, abbreviated as “c5-cre22”). Normal 46,XY karyotype is shown.

D) Immunolabelling of BU3 NGPT c5-cre22 iPSCs for pluripotency markers (NANOG, SSEA4, SOX2, OCT4 and TRA1-81) and DAPI to counterstain nuclei (scale bar=100μm). Phase microscopy of a representative BU3 NGPT iPSC colony (scale bar=400μm).

E) Immunolabeling of BU3 NGPT iPSCs for classic endoderm (SOX17), primitive streak (Brachyury) and ectdodermal markers (TUJ1, NESTIN). Nuclei are stained with DAPI. Scale bar=100μm.

### Supplemental Figure 2:

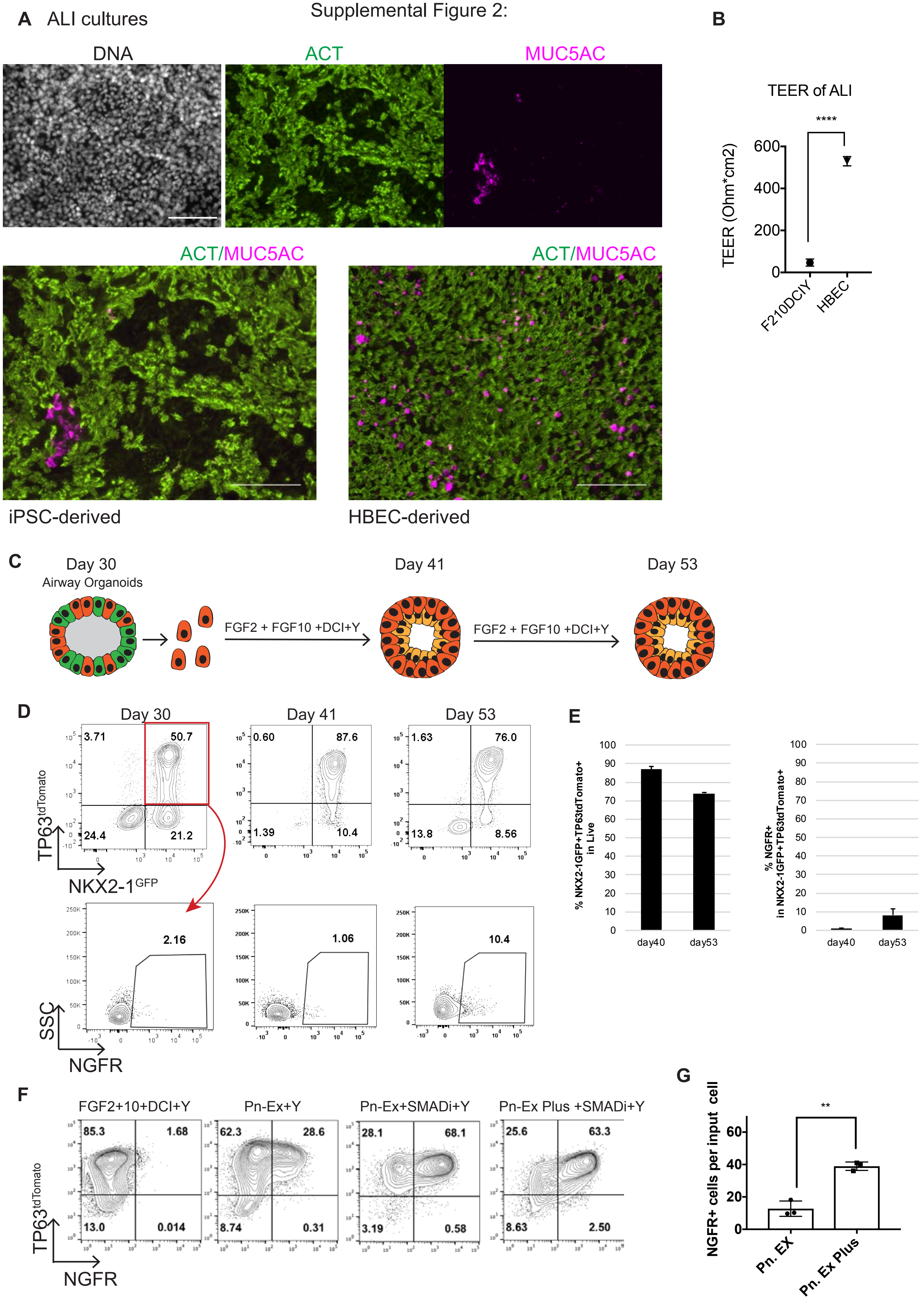

**Supplemental figure 2: Assessment of the differentiation capacity and NGFR expression of NKX2-1$^{GFP+}$;TP63$^{tdTomato+}$ cells in FGF2+10+DCI+Y medium.**

A) Representative images of ALI cultures generated from sorted GFP+/TOM+ cells (day 40-42) in FGF2+10+DCI+Y medium immunolabelled with antibodies against ACT and MUC5AC and with nuclear labelling using Hoechst (upper panel). Merge images iPSC-derived ALI compared to primary HBEC-derived ALI (lower panel). Scale bars=100μm.

B) TEER of iPSC-derived ALI (n=4) from (A) compared to HBEC-derived ALI (n=4). (****=p<0.0001)

C) Schematic of experiment. GFP+/TOM+ cells were sorted on day 30 of differentiation and cultured in FGF2+10+DCI+Y medium until day 53.

D) Representative flow cytometry plots from the experiment depicted in (C) analyzing the expression of GFP, tdTomato and NGFR on days 30, 41 and 53.

E) Quantification of the percent GFP+/TOM+ cells on days 41 (87.0±1.5, mean±SD, n=3) and 53 (73.8±0.7) of the experiment summarized in (C) and the frequency of NGFR+ cells within GFP+/TOM+ population (days 41=0.8±0.3 and day 53=8.0±3.3).

F) On day 35 of differentiation (see Figure 2A schematic) GFP+/TOM+ cells were sorted and plated in continued FGF2+10+DCI+Y medium, PneumaCult-EX medium supplemented with Y-27632 (Pn-Ex+Y), PneumaCult-EX medium supplemented with A83-01, DMH-1, and Y-27632 (Pn-Ex+SMADi+Y), or PneumaCult-EX Plus medium supplemented with A83-01, DMH-1, and Y-27632 (Pn-Ex Plus+SMADi+Y) until day 50. Representative flow cytometry plots of GFP and tdTomato expression are shown.

G) NGFR+ cell on day 50 per input cell GFP+/TOM+ cell on day 35 of differentiation in PneumaCult-EX medium supplemented with A83-01, DMH-1, and Y-27632 (Pn-Ex), or PneumaCult-EX Plus medium supplemented with A83-01, DMH-1, and Y-27632 (Pn-Ex Plus).

Supplementary Figure 3: ScRNA-Seq of BU3 NGPT iPSC in FGF2+10+DCI+Y vs BC media.

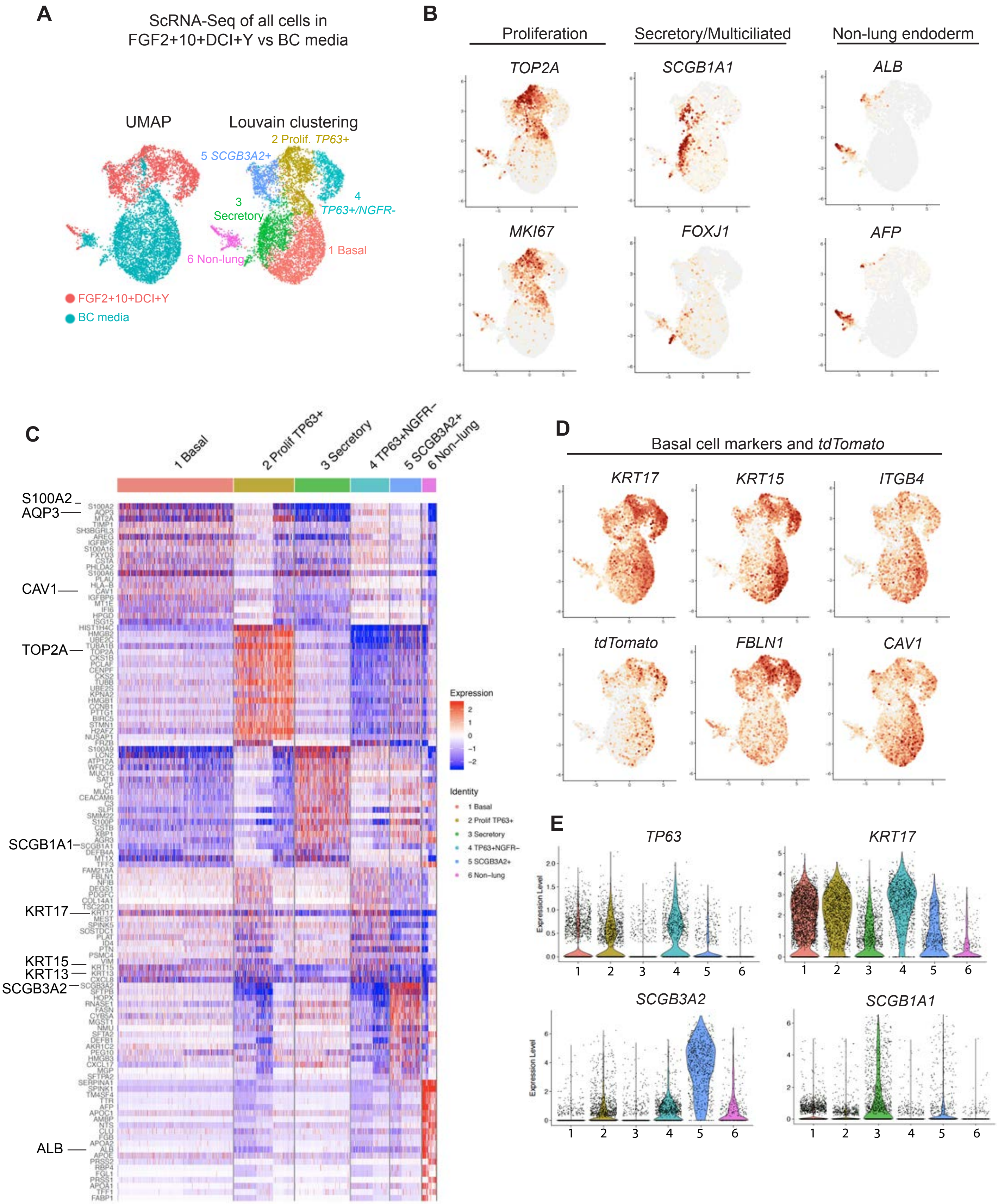

**Supplemental figure 3: Sc-RNA-Seq analysis of airway epithelial organoids in FGF2+FGF10+DCI+Y or BC media.**

A) ScRNA-Seq of day 45 cells from primary BC or FGF2+FGF10+DCI+Y media (identical to Figure 2E). UMAP (left panel) displays the culture conditions used to treat cells. Louvain clustering (res = 0.25) identifies 6 clusters (1-6). See also Figures 2E-H.

B) UMAPs of key markers used to classify the identity of cell clusters including proliferation (*TOP2A* and *MKI67*), secretory (*SCGB1A1*) and multiciliated (*FOXJ1*) cells and non-lung endoderm (*ALB* and *AFP*).

C) Gene-expression heatmap of the top 20 DEGs across clusters 1-6. Selected canonical lineage and proliferation markers for each cluster are highlighted.

D) UMAPs of the expression pattern of additional basal cell markers and tdTomato. See also Figure 2F.

E) Gene expression violin plots of additional secretory and basal markers across clusters 1-6.

Supplementary Figure 4: scRNA-Seq of primary BCs and ALI cultures

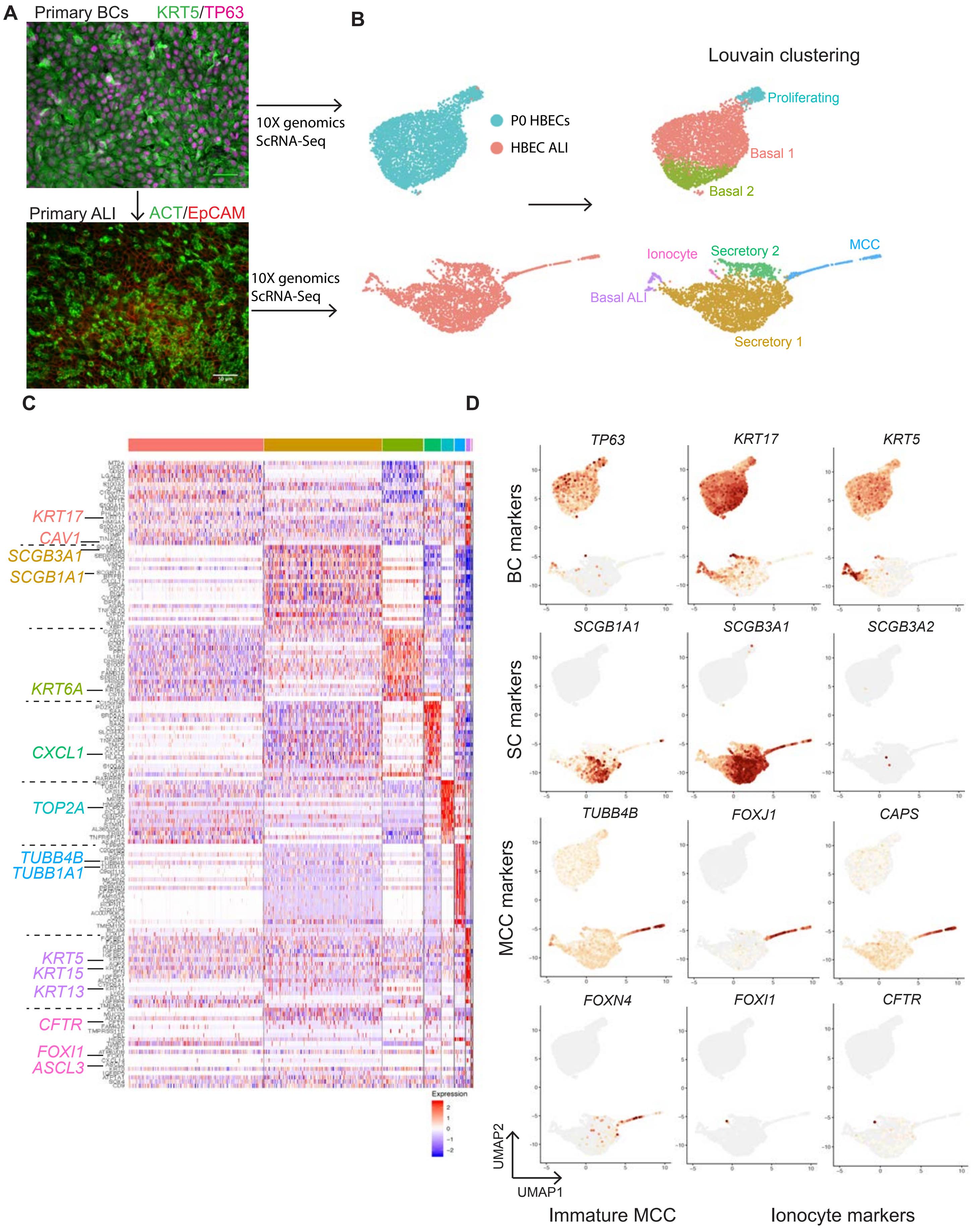

**Supplemental figure 4: Sc-RNA-Seq of primary HBECs and their differentiated progeny.**

A) Schematic of scRNA-Seq experiment profiling P0 primary HBECs (DD001m) in BEGM medium and after 20 days of differentiation into a mucociliary epithelium in ALI culture. These data were used to generate the gene-signatures for BC, SC, MCC and ionocytes (Table 1) applied to iBCs (Figure 2) and iBC-derived ALI (Figure 4). From this scRNA-Seq experiment we performed immunolabelling of DD001m HBEC cultures with antibodies against basal cell markers TP63 and KRT5 (scale bar=50μm) and ALI cultures with antibodies against ACT and EpCAM (scale bar=100μm).

B) UMAP of cells from P0 and ALI conditions after filtering out doublets and degraded cells. 3,943 cells from P0 HBECs and 3160 cells from ALI culture were analyzed. UMAPs labelled with original identity (left graph) and after Louvain clustering and annotation of clusters (right panel) are shown.

C) Gene-expression heatmap of the top 20 DEGs across 8 clusters. Select canonical lineage and proliferation markers for each cluster are highlighted.

D) UMAPs of the gene-expression of a panel of canonical markers of basal, secretory, immature multiciliated, multiciliated cells and ionocytes.

Supplementary Figure 5: Expansion, cryopreservation and differentation of iBCs

**A** Extended passage of iBCs

BU3 NGPT 9 passages after GFP+/TOM+ sort

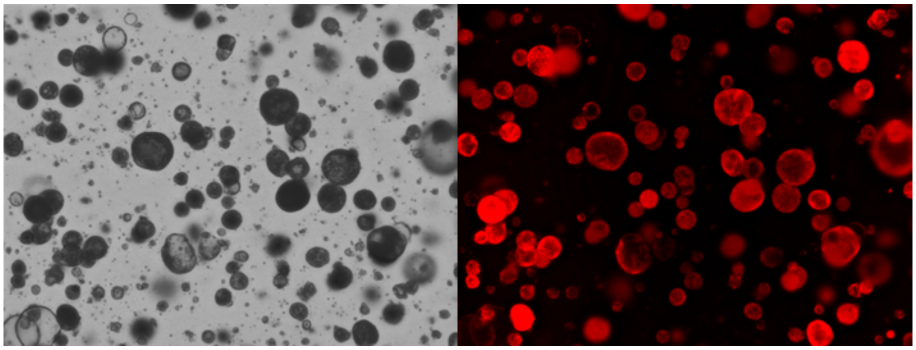

**B**

BU3 NGPT Karyotype

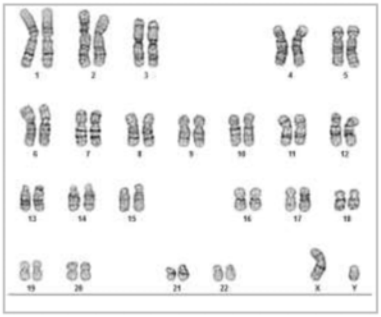

**C** Limiting dilution assay

BFP/GFP/dsRed

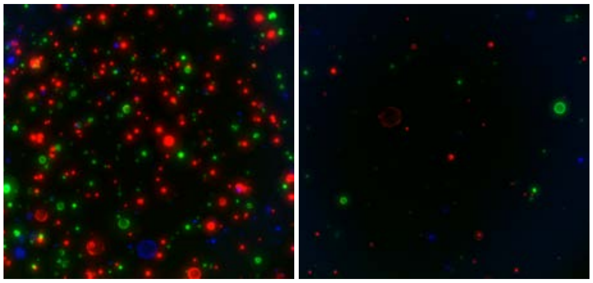

1000 cells/ul 200 cells/ul

Percent clonal spheres

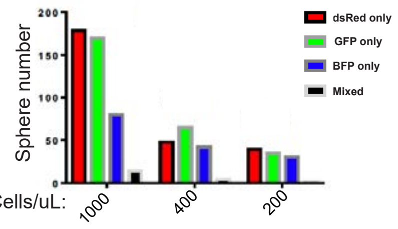


Clonal(%): 95.7 97.5 99.1

ACT/DNA

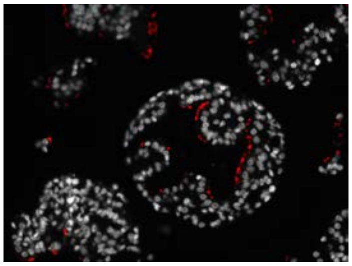

**D** Multi-lineage differentiation in ALI after extended passaging of iBCs

DNA ACT/MUC5AC

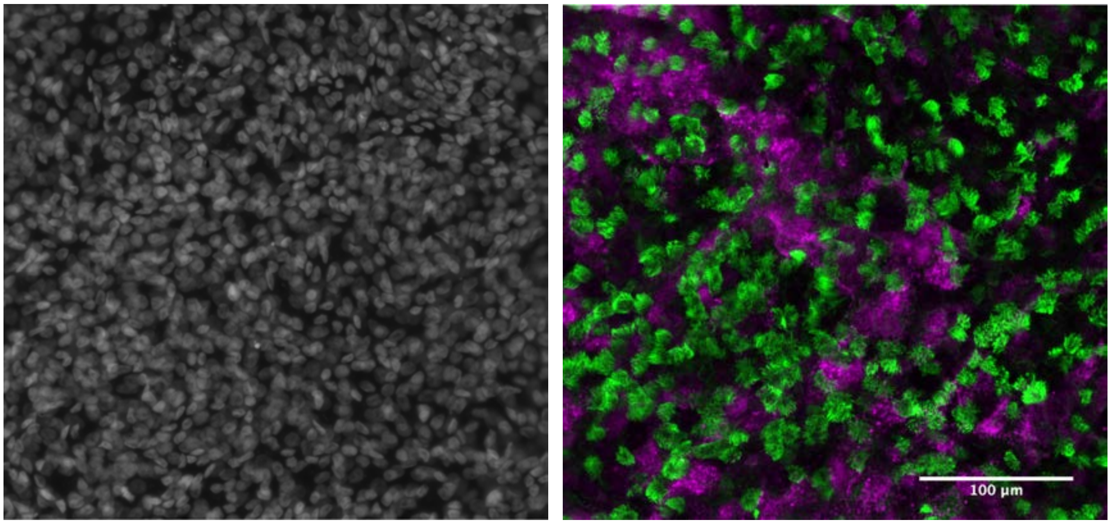


ALI after 10 passages of iBCs

**Supplemental figure 5: Expansion, cryopreservation and multi-lineage differentiation of iBCs.**

A) Representative images, phase (left) and fluorescence (right), of iBCs after 9 passages in 3D Matrigel culture.

B) Normal karyotype (46,XY) of BU3 NGPT iPSC-derived iBCs on day 82 of directed differentiation.

C) Representative images of 1000 cells/μL and 200 cell/μl concentrations from the limiting dilution assay used for plating density to allow clonal outgrowth of spheres (see Figure 3C). iPSC-derived airway progenitors were infected with one of three lentiviral vectors designed to constitutively express either GFP, BFP or dsRed. Transduced cells were sorted, combined and replated in 3D Matrigel at a range of concentrations. The number of clonal and non-clonal spheres at each density was quantified (middle graph). Clonal spheres from iBCs cultured in ALI differentiation medium were immunolabelled with an antibody against ACT and nuclei labelled with Hoechst.

D) ALI cultures from BU3 NGPT derived iBCs after 10 passages in 3D culture immunolabelled with antibodies against ACT and MUC5AC (nuclei stained with Hoechst; scale bar=100μm).

Supplementary Figure 6: ScRNA Seq of iPSC-derived ALI

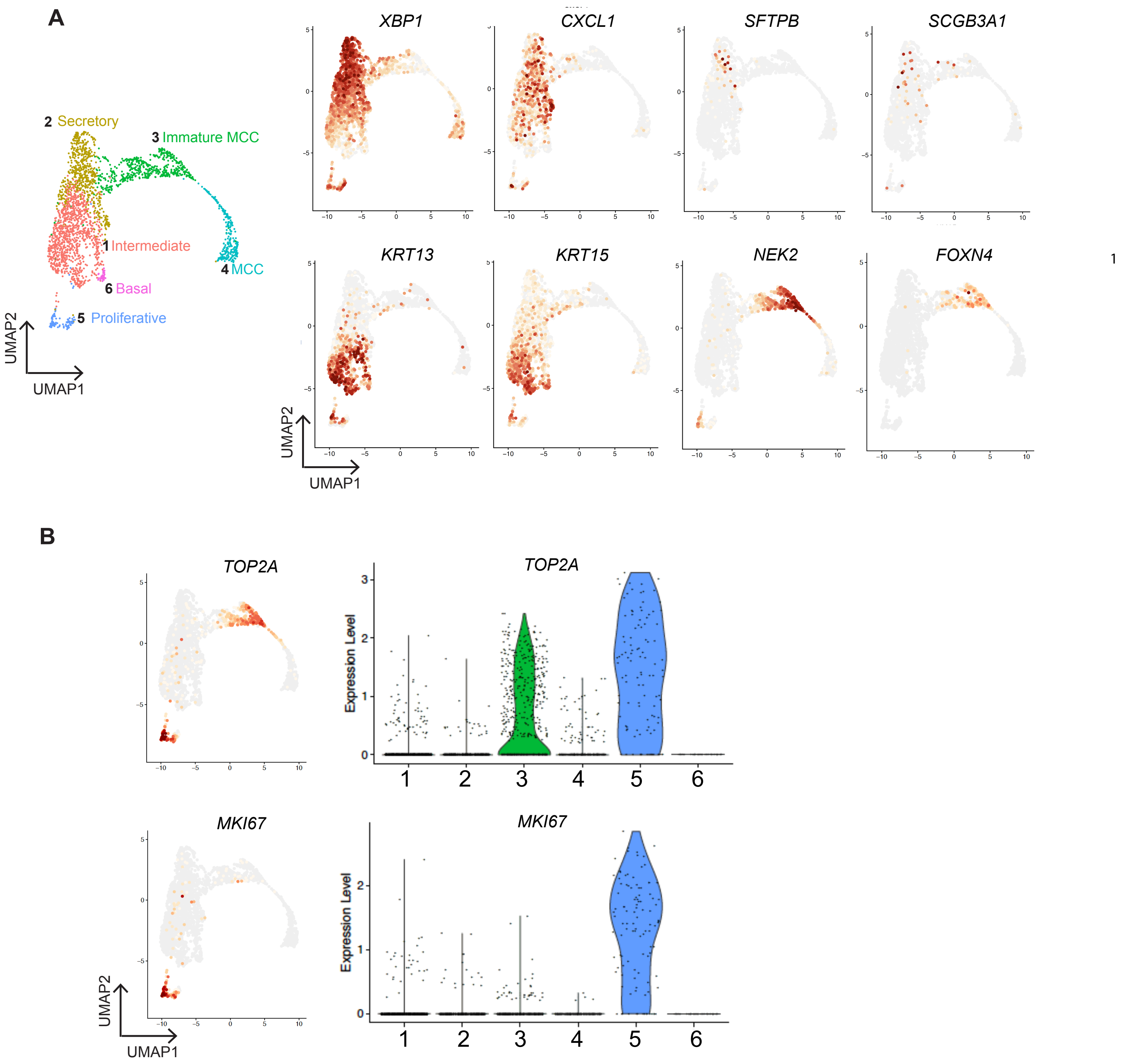

**Supplemental figure 6: Sc-RNA-Seq of iBC-derived ALI.**

A) UMAP of iBC-derived ALI with Louvain clustering (clusters 1-6) and annotation (identical to Figure 4A). The expression of additional genes identified as highly differentially expressed genes within in Figure 4B are presented. See also Figure 4E.

B) UMAP and violin plots of the gene expression of cell cycle genes *TOP2A* and *MKI67*.

Supplementary Figure 7: Disease modeling using iBCs

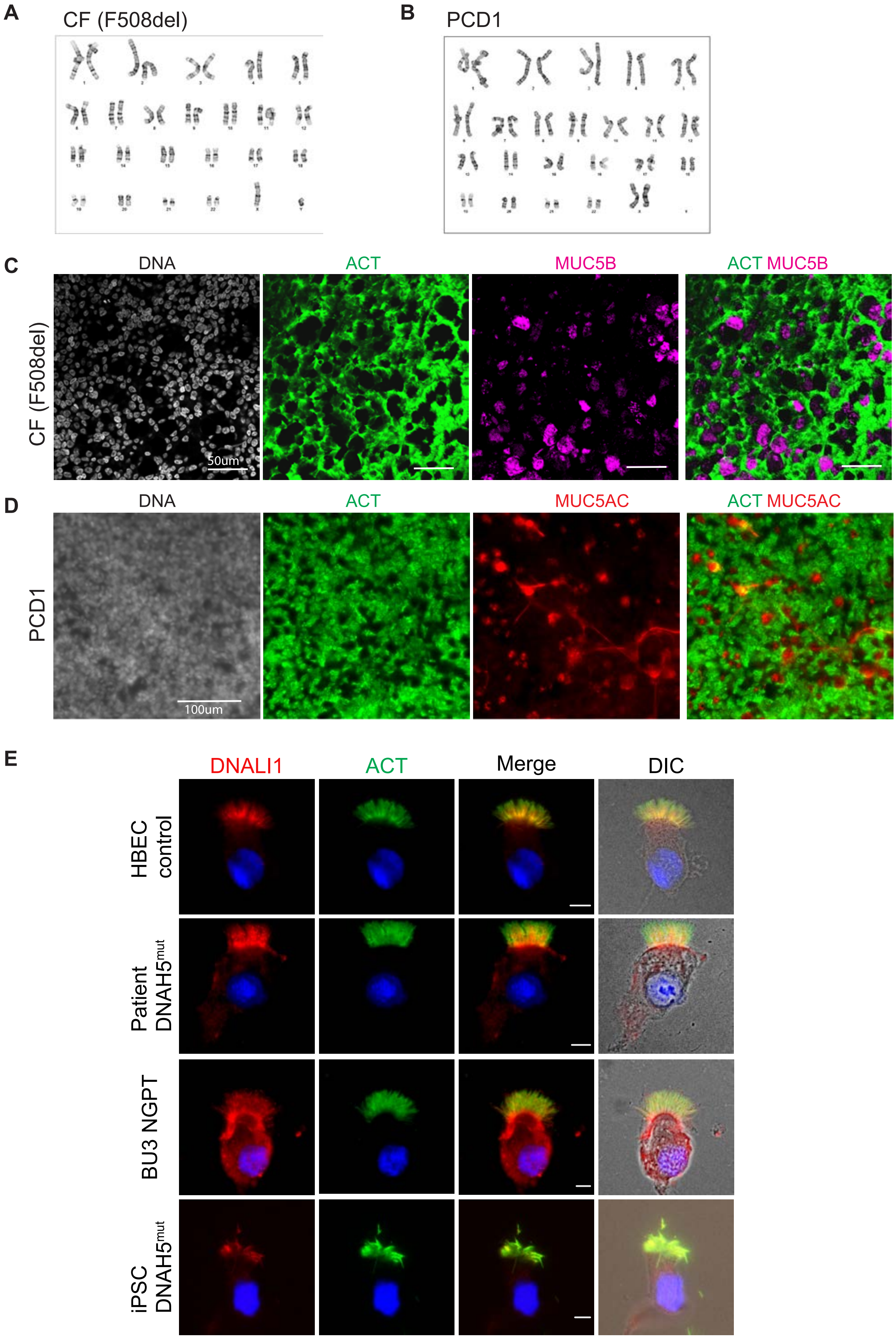

**Supplemental figure 7: Characterization of CF and PCD iPSCs and their airway epithelial derivatives.**

A) G-band karyotyping of F508del iPSCs demonstrating a normal 46,XY karyotype. B) G-band karyotyping of mutant DNAH5 iPSCs demonstrating a normal 46,XX karyotype.

C) Characterization of mucociliary differentiation with antibodies against ACT and MUC5B in ALI cultures generated from F508del iPSCs (using the protocol summarized in Figure 6A) in the experiments described in Figures 7B-C. Nuclei stained with Hoechst. Scale bar=50μm.

D) Characterization of mucociliary differentiation with antibodies against ACT and MUC5AC in ALI cultures generated from *DNAH5* mutant iPSCs in the experiments described in Figures 7D-G. Nuclei stained with Hoechst. Scale bar=100μm.

E) Immunolabeling of multiciliated cells from the samples detailed in Figure 7D with antibodies against ACT and DNALI1 (nuclei stained with DAPI; scale bar=10μm).

**Table 1**

| | Basal | Secretory | Multiciliated | Ionocyte |
|---|---|---|---|---|
| 1 | *UPP1* | *SCGB3A1* | *TPPP3* | *TMEM61* |
| 2 | *MT2A* | *WFDC2* | *C20orf85* | *CRYM* |
| 3 | *LGALS1* | *MSMB* | *CAPS* | *MUC20* |
| 4 | *S100A2* | *BPIFB1* | *RSPH1* | *ANXA4* |
| 5 | *AREG* | *SERPINB3* | *TUBB4B* | *CFTR* |
| 6 | *ZFAS1* | *VMO1* | *TUBA1A* | *FAM43A* |
| 7 | *G0S2* | *SLPI* | *C9orf116* | *TMPRSS11E* |
| 8 | *S100A14* | *BPIFA1* | *PIFO* | *ITPR2* |
| 9 | *S100A10* | *CD74* | *MORN2* | *CEL* |
| 10 | *KRT17* | *CXCL17* | *C5orf49* | *NREP* |
| 11 | *KRT6A* | *PIGR* | *PSENEN* | *MUC20-OT1* |
| 12 | *HMGA1* | *SCGB1A1* | *CFAP126* | *ATP6V0A4* |
| 13 | *PHLDA1* | *CXCL1* | *FAM183A* | *AKR1B1* |
| 14 | *TMSB10* | *PSCA* | *FOXJ1* | *HES6* |
| 15 | *TIMP1* | *AGR2* | *AGR3* | *TIMP3* |
| 16 | *ADIRF* | *HLA-DRA* | *FAM229B* | *LRMP* |
| 17 | *TINAGL1* | *TNFSF10* | *KIF9* | *PLCG2* |
| 18 | *S100A16* | *CYP2F1* | *DNALI1* | *GADD45G* |
| 19 | *IGFBP6* | *GLUL* | *LRRIQ1* | *KIT* |
| 20 | *SH3BGRL3* | *XBP1* | *DYNLL1* | *AZGP1* |
| 21 | *C16orf74* | *STATH* | *IFT57* | *ATP6V0B* |
| 22 | *SNHG8* | *LYPD2* | *ODF3B* | *HEPACAM2* |
| 23 | *RPS21* | *S100A9* | *IFT22* | *ASCL2* |
| 24 | *RPL37A* | *GSN* | *NUDC* | *MGLL* |
| 25 | *RPL37* | *HLA-DRB5* | *CETN2* | *SDC2* |
| 26 | *IER3* | *ALDH1A1* | *UFC1* | *SIRT2* |
| 27 | *LAMC2* | *C3* | *TCTEX1D2* | *FOXI1* |
| 28 | *ITGB1* | *RARRES1* | *CFAP298* | *MYO6* |
| 29 | *EIF4EBP1* | *LY6E* | *DPCD* | *IQGAP2* |
| 30 | *RPL36* | *RARRES3* | *WDR54* | *DMRT2* |

**Table 1:** Gene-expression signatures of the top 30 DEGs in human BCs, SCs, MCCs and ionocytes based on ScRNA-Seq profiling of primary P0 HBECs and after differentiation in ALI culture. See also Figure 2G, 4B and S3.

**Supplemental Video 1: En-face video of iBCs (BU3 NGPT) differentiated in ALI conditions.**

**Supplemental Videos 2A-D: Representative high magnification videos of the samples analyzed in Figure 7D to measure ciliary beat frequency.**

**Methods:**

#### Human subjects

The Institutional Review Board of Boston University approved the generation and differentiation of human iPSCs with documented informed consent obtained from participants. The generation of CF iPSCs was approved by Boston Children's Hospital Institutional Review Board. Human airway tissue and primary HBECs were received from the CF Center Tissue Procurement and Cell Culture Core (Dr. Scott Randell) at the University of North Carolina. Human lung tissue was procured under the University of North Carolina Office of Research Ethics Biomedical Institutional Review Board protocol No. 03-1396. For PCD studies, human protocols were approved by the institutional review board at Washington University in St. Louis. Subjects with known mutations causative of PCD were recruited from the PCD and Rare Airway Disease clinic at St. Louis Children's Hospital and Washington University. Informed consent was obtained from individuals (or their legal guardians). All human samples were de-identified.

#### Human iPSC reprogramming and iPSC/ESC maintenance

All iPSC and ESC lines were maintained in feeder-free conditions on hESC-qualified Matrigel (Corning) in StemFlex Medium (Thermo Fisher Scientific, Waltham, MA) or mTeSR1 medium (Stemcell Technologies, Vancouver, Canada) and passaged with Gentle Cell Dissociation Reagent (Stemcell Technologies) or ReLeSR (Stemcell Technologies). All human ESC/iPSC lines used were characterized for pluripotency and were found to be karyotypically normal. The reprogramming and gene-editing methods to derive BU3 iPSCs and target the NKX2-1 locus to generate BU3 NKX2-1$^{GFP}$ iPSCs were previously described (Hawkins et al. 2017). PCD1 iPSC line (*DNAH5* mutation) was generated by reprogramming peripheral blood mononuclear cells with the human EF1a-STEMCCA-loxp lentiviral vector followed by Cre-mediated vector excision, according to our detailed protocol (Sommer et al. 2012). DD001m iPSCs were generated from "DD001m" P0 primary HBECs by modifying a Sendai virus based reprogramming protocol to substitute the base medium for bronchial epithelial growth medium (Park & Mostoslavsky 2018). Homozygous F508del (p.Phe508del) iPSCs were generated from peripheral blood mononuclear cells at Boston Children's Hospital Stem Cell Program by Sendai virus using the Cytotune 2 kit (Thermo Fisher), yielding clones 791 and 792 ("CF"). Clone 792 was selected for correction of the p.Phe508del deletion and the experiments in Figure 7. The RUES2 human embryonic stem cell line was a generous gift from Dr. Ali H. Brivanlou of The Rockefeller University, New York City, NY. Standard immunolabeling for pluripotency markers and G-band karyotyping was performed to ensure pluripotency and normal karyotype in clones selected for these experiments (as shown in Figure S1). For BU3 NGPT iPSCs spontaneous differentiation into mesoderm, endoderm and ectodermal lineages was demonstrated by generating embryoid bodies with iPSC medium in ultra low attachment plate (Corning) and then replacing with base medium supplemented with 10% fetal bovine serum in adherent culture. The cell types from each lineage was identified with standard immunolabeling (as shown in Figure S1) using anti-bodies listed in key resources table. Primocin (Invivogen, San Diego, CA) was routinely added to prevent mycoplasma contamination and all iPSC and ESC lines screened negative for mycoplasma contamination and were routinely tested and remained negative.

**Basal cell reporter iPSC line generation**

The dual reporter, NKX2-1$^{GFP}$ and P63$^{tdTomato}$, iPSC lines (BU3 NGPT) were derived from the published single reporter, NKX2-1$^{GFP}$, iPSC line (BU3 NG), a normal donor iPSC carrying homozygous NKX2-1$^{GFP}$ reporters (Hawkins et al. 2017). The BU3 NG line was targeted and integrated with a P63$^{tdTomato}$ fluorescent reporter using CRISPR/Cas9 technology. A gRNA was designed to target exon 4 of the endogenous *TP63* (target sequence TGCGCGTGGTCTGTGTTATA) and a donor template was constructed to contain the tdTomato sequence and removable antibiotic selection cassette via Cre recombination flanked by arms of homology. In this report we used the clone BU3 NGPT c5-cre22, where P63$^{tdTomato}$ fluorescent reporter-integrated clone 5 was further cre-excised to remove selection cassette (sub-clone 22). **Monoallelic correction using CRISPR/Cas9 editing of** F508del homozygous **iPSCs**

CRISPR gene repair was performed on clone 792 by nucleofecting 1M TrypLE-Select dissociated cells with 5µg Cas9-GFP plasmid (Addgene, 44719), 5 µg of a U6 promoter driven sgRNA plasmid (target sequence ACCATTAAAGAAAATATCAT), 10µg of a plasmid containing a 1.4kb piece of the *WT CFTR* that includes the exon encoding F508. Repaired clones were identified by ddPCR and Sanger sequencing of long-range PCR products. Clones 1566 (CFTR WT/ F508del) and 1567 (CFTR F508del / F508del) were confirmed by DNA STR fingerprint (match with donor) and karyotype analysis (no abnormalities).

**Directed Differentiation of hPSCs into airway organoids**

This protocol is based on previously described approaches to derive airway organoid from hPSCs (Hawkins et al. 2017; McCauley et al. 2017) including a detailed step-by-step protocol prior to basal cell medium (McCauley et al. 2018). NKX2-1+ lung progenitors were generated from hPSCs first by inducing definitive endoderm with STEMdiff Definitive Endoderm Kit (Stemcell Technologies) for 60-72 hours. Endoderm-stage cells were dissociated and passaged in small clumps to hES-qualified Matrigel-coated (Corning) tissue culture plates (Corning) in a complete serum free differentiation medium (cSFDM) consisting of a base medium of IMDM (Thermo Fisher, Waltham, MA) and Ham's F12 (Thermo Fisher) with B27 Supplement with retinoic acid (Invitrogen, Waltham, MA), N2 Supplement (Invitrogen), 0.1% bovine serum albumin Fraction V (Invitrogen), monothioglycerol (Sigma, St. Louis, MO), Glutamax (Thermo Fisher), ascorbic acid (Sigma), and antibiotics. To pattern endoderm into anterior foregut the cSFMD base medium was supplemented with 10 μM SB431542 (Tocris, Bristol, United Kingdom) and 2 μM Dorsomorphin (Stemgent, Lexington, MA) for 72 hours (Ref, green et al). Cells were then cultured for 9-11 additional days (typically, 144 hr - day 15) in cSFDM containing 3 μM CHIR99021 (Tocris), 10 ng/mL recombinant human BMP4 (rhBMP4, R&D Systems), and 50-100 nM retinoid acid (Millipore-Sigma) to induce a NKX2-1 primordial lung progenitors(Serra et al. 2017). NKX2-1+ lung progenitors were sorted by NKX2-1$^{GFP}$ expression for BU3 NGPT or enriched based on the expression of cell surface markers CD47hi/CD26- for non-reporter iPSC (see below) and then were resuspended in three-dimensional growth factor reduced Matrigel at a density of 400 cells per μl and pipetted in 25-50 µl droplets onto the base of tissue culture plates (Hawkins et al. 2017). After 10-15 minutes at 37°C to allow gelling of Matrigel airway medium composed of cSFDM containing 250ng/ml FGF2 (rhFGFbasic; R&D Systems), 100ng/ml FGF10, 50nM dexamethasone, 100nM 8-Bromoadenosine 3′,5′-cyclic monophosphate sodium salt (cAMP; Millipore-Sigma), 100nM 3-Isobutyl-1-methylxanthine (IBMX; Millipore-Sigma), and 10µM Y-27632 (Y; Tocris) (FGF2+10+DCI+Y) was added.

**Purification of NKX2-1+ Lung Progenitors, airway progenitors of iBCs by Cell Sorting**

This sorting protocol is further detailed in previous publications (Hawkins et al. 2017; McCauley et al. 2018). On ~ day 15, cells were harvested by incubation with 0.05% Trypsin-EDTA for 15-20 min at 37C. Trypsin was neutralized and cells were washed with medium containing 10% fetal bovine serum (FBS, Thermo Fisher.) Harvested cells were spun at 300 RCF for 5 min and resuspended in the buffer containing Hank's Balanced Salt Solution (Thermo Fisher), 2% FBS, 25mM HEPES 2mM EDTA (FACS buffer) supplemented with 10µM Y-27632 and stained with propidium iodide (PI, ThermoFisher) or calcein blue AM (Thermo Fisher) for live cell selection during flow cytometry. Live cells were sorted based on NKX2-1$^{GFP}$ expression for BU3 NGPT or by staining for CD47 (Biolegend) and CD26 (Biolegend) for non-reporter PSCs and gating for CD47hi/CD26-. Airway progenitors were sorted on ~ day 30 of directed differentiation. Organoids cultured in 3D Matrigel were harvested by incubation with 1 U/mL Dispase (Stemcell technologies) or 2mg/ml Dispase (Thermo Fisher) for ~ 60 min at 37C until Matrigel was fully dissolved. Dissociated organoids were collected with a 1000µl pipette, transferred to a 15ml conical and and centrifuged at 200 RCF for 3 min. The cells were resuspended and incubated in 0.05% trypsin at 37°C for approximately 10min until a single cell suspension was achieved. Trypsin was inactivated by adding 10% fetal bovine serum (Hyclone) in DMEM (Gibco). Cells were then centrifuged at 300 RCF for 5 min at 4°C, counted and resuspended in FACS buffer. 10μM of the cell viability dye Calcein blue (ThermoFisher) was added and Calcein Blue+/NKX2-1$^{GFP+}$/TP63$^{tdTomato+}$ or PI$^{-}$/NKX2.1$^{GFP+}$/TP63$^{tdTomato+}$ cells were sorted and replated at 400 cells per μl of density in Matrigel (Corning) and cultured in FGF2+10+DCI+Y medium. One day after (unless indicated), culture medium was switched to Pneumacult ExPlus (Stemcell Technologies) supplemented with 1μM A83-01 (Tocris), 1μM DMH1 (Tocris), and 10μM Y-27632 (Basal cell medium). In some cases, Pneumacult Ex (Stemcell technologies) supplemented with 1μM A83-01 (Tocris), 1μM DMH1 (Tocris), and 10μM Y-27632 was used as indicated in figures. For non-reporter iPSCs, single cells dissociated from airway organoids in FGF2+10+DCI+Y were replated without a sorting step and changed to basal cell medium as above. Viable iBSCs at ~ Day 40 of directed differentiation or later were sorted based by dissociating organoids to a single-cell suspensions as above and flow sorting based NKX2-1GFP+/TP63tdTomato+/NGFR+ cells for BU3 NGPT, or by labelling with mouse monoclonal anti-NGFR (Cat.# 345108, Biolegend) or mouse monoclonal anti-EpCAM (Cat.# 24234, Biolegend) with isotype controls (mouse monoclonal IgG1k APC-conjugated, Cat.#400122, Biolegend). Both anti-NGFR and anti-EpCAM antibodies were used at a dilution of 1:100 in a suspension of 1x10$^{6}$ cells per 50 or 100μL on ice for 30 min. For these sorting experiments a MoFlo Astrios Cell sorter (Beckman Coulter, Indianapolis, IN) was used at Boston University Medical Center Flow Cytometry Core

Facility or BD FACSMelody (BD biosciences) was used at the UTHealth Flow Cytometry Service Center. Sorted iBCs were either replated in ALI culture (see below) or resuspended in Matrigel (Corning) droplets at 400 cells/ul, plated and cultured in basal cell medium as above. iBCs in 3D Matrigel (Corning) droplets were fed with basal cell medium every 2-3 days and required passaging every ~ 10-14 days. iBCs were cryopreserved by first dissociating organoids to single-cells as above and resuspending in basal cell medium supplemented with 10% dimethylsulfoxide (ThermoFisher) and transferred to cryovials. Cryovials were placed in cryostorage containers in a -80°C freezer to cool at -1°C /min for 24 hours and then transferred to and -150°C freezer the following day for long-term storage. iBCs were thawed by warming to 37°C in a water or bead bath, diluting in 5-10ml of basal cell medium (37°C) added dropwise and centrifuged at 300 RCF for 5min. The cell pellet was then resuspended in Matrigel (Corning) at 1000cells/µl, plated in tissue culture plates and after 10-15 min at 37°C basal medium was added to the well.

**Air Liquid Interface culture of iBCs or other iPSCs-derived airway epithelial cells**

To differentiate iBCs in ALI culture we adapted existing, detailed protocols for primary HBECs (Fulcher & Randell 2013). iPSCs-derived airway epithelial cells including iBCs were seeded on 6.5 mm Transwells with 0.4 µm pore polyester membrane inserts (Corning Inc., Corning, NY), coated with Matrigel (Corning) diluted in DMEM/F12 (Gibco) according to the recommended dilution factor from the lot-specific certificate of analysis, at 200,000 cells per insert with a medium used to culture each cell culture (e.g. iBCs cultured in Basal cell medium were plated on Transwells and cultured in Basal cell medium) for 3 to 7 days. Once visual inspection confirmed the Transwell membranes were covered by a confluent sheet of cells the culture medium was switched to PneumaCult-ALI medium (Stemcell Technologies) in both apical and basal chambers. The following day, the medium from top chamber was removed and the cells were further differentiated in air-liquid interface conditions for ~ 2 weeks or longer before analysis.

**Primary human bronchial epithelial cells and human airway tissue.**

De-identified, cryopreserved HBECs were received from the CF Center Tissue Procurement and Cell Culture Core, Marsico Lung Institute, University of North Carolina where they were harvested and cultured as previously described in detail. Paraffin-embedded sections of de-identified human airway tissue were also received. These samples are exempt from regulation by HHS regulation 45 CFR Part 46. Freshly isolated P0 HBECs were expanded in culture on collagen type I/III-coated tissue culture plates in non-proprietary bronchial epithelial growth medium (BEGM). At 70-90% confluence HBECs were dissociated with Accutase (Sigma), counted and transferred to human placental type IV collagen-coated membranes (Transwell, Corning Inc., Corning, NY) at a density of $1 \times 10^5$ cells per $cm^2$ and differentiated using air-liquid interface conditions in a non-proprietary medium, “UNC-ALI” medium for at least 20 days before analysis.

**Flow cytometry analysis**

Flow cytometry analysis was performed as described above. NKX2-1$^{GFP}$, TP63$^{tdTomato}$ expression or fluorochrome-conjugated primary antibodies (listed in the Key Resources Table) were detected using a FACS Calibur (BD Biosciences), Stratedigm S1000 EXi

(Stratedigm Inc, San Jose, CA) or MoFlo Astrios (Beckman Coulter) at Boston University Medical Center or with BD FACSMelody and BD FACSChorus software (BD Biosciences) at the UTHealth Flow Cytometry Service Center. Calcein blue or PI was used to identify viable cells depending on the flow cytometer used. FlowJo software (BD biosciences) was used for further analysis.

**Airway epithelial cell immunofluorescence staining**

Organoids in 3D Matrigel were harvested by incubation in Dispase as described above, and then fixed in 4% paraformaldehyde (PFA) (ThermoFisher or Electron Microscopy Sciences, Hatfield, PA) for four hours at room temperature, paraffin-embedded and immunostained as previously described (McCauley et al. 2018; McCauley et al. 2017). ALI cultures were fixed by placing Transwell filters in 4% PFA at 4°C overnight. Either whole mount immunostaining of Transwell filters or paraffin-embedding was performed. For paraffin sections, fixed samples were dehydrated with a series of increasing concentrations of ethanol, cleared with xylene and infiltrated with paraffin before embedding in wax. Sections of 5 µm thickness were transversely cut using an HM 325 Rotary Microtome (Thermo Fisher Scientific). Sectioned samples were stained with hematoxylin and eosin. For immunofluorescence analysis, antigen retrieval was performed on sectioned samples using Antigen retrieval reagent-Basic (R&D Systems) or Antigen Unmasking Solution, Citric Acid Based (Vector Labs, Burlingame CA) following manufacturer’s instructions. Whole inserts and sectioned samples were stained with primary and secondary antibodies (antibody information and sources are detailed in the Key Resources Table). Briefly, samples were permeabilized with with 0.1% or 0.3 %

Triton X-100 (Sigma) in PBS for 15-30 min and blocked with 2 % bovine serum albumin (BSA) with or without 4% Normal Donkey Serum and 0.1% Triton X-100 for 1 hour. Samples were then incubated with primary antibodies overnight at 4 ºC, followed by the incubation with the respective secondary antibodies at room temperature for one to two hours. Prolong™ Gold Antifade Mountant with DAPI (Thermo Fisher Scientific) was added to counter-stain and coverslipped. Images were acquired using a Leica DMi8 microscope (Leica Microsystems, Wetzlar, Germany) and Leica Application Suite Software (Leica Microsystems) or a Nikon Eclipse Ni-E microscope (Melville, NY) . Confocal microscopy was performed for whole mount staining sample stained with antibodies as described above and counter-stained with DRAQ5 (Cell Signaling Technology, Beverely, MA) using Leica Application Suite Software.

To characterize the ciliary defect in *DNAH5* mutant cells (Figure 7 and supplemental videos), airway cells were fixed and immunostained as previously described [2, 4]. Primary and secondary antibodies used to detect dynein arms included anti-DNAH5 (1:100, HPA037470, Millipore-Sigma, St. Louis, MO), ACT (1:4000, clone 6-11-B1, Millipore-Sigma), DNALI1 (1:100, HPA028305, Millipore-Sigma). Primary antibodies were detected using fluorescently labeled secondary antibodies (Alexa Fluor, Life Technologies, Grand Island, NY, USA). Nuclei were stained using 4’, 6-diamidino-2-phenylindole (DAPI) (Vector Laboratories, Burlingame, CA, USA) or Images were acquired using an epiflourescent microscope interfaced with imaging software (LAS X, Leica, Buffalo Grove, IL) and adjusted globally for brightness and contrast using Affinity Photo (Serif Ltd, Nottingham, UK).

**Single-cell RNA-sequencing**

Cells were prepared for sc-RNA-Seq by dissociating organoids or ALI cultures using the methods described above. Live cells were sorted on a MoFlo Astrios Cell sorter (Beckman Coulter, Indianapolis, IN) at Boston University Medical Center Flow Cytometry Core and scRNA-Seq was performed using the Chromium Single Cell 3' system (10X Genomics) at the Single Cell Sequencing Core at Boston University Medical Center according to the manufacturer's instructions (10X Genomics). Facility Single cell reads were aligned to the reference genome GRCh38 to obtain a gene-to-cell count matrix with Cell Ranger version 3.0.2 (10x Genomics). The average sample had a mean of 40,154 reads per cell (ranging from 18,415 to 50,219 depending of the library). The median number of genes detected per cell on the average sample was 3,303 genes. This matrix was pre-processed using Seurat version 3.1.0 (Butler et al. 2018), and filtered to remove stressed or dead cells (those with a high percentage of reads mapping to mitochondrial genes, that is beyond 10-20%, depending on the shape of the distribution of mitochondrial reads per cell) and potential doublets (where the number of genes detected is above a percentile set as threshold based on the expected proportion of doublets at any given cell density (100 - (number of cells/1000) / 100). After normalizing, scaling and regressing out unwanted sources of variation (like cell degradation), we identified highly variable genes and used them for linear dimensionality reduction (PCA). The first 20 principal components were then used for clustering based on the Louvain method for community detection at different resolutions (from 0.25 to 1.5 at intervals of 0.25). Further non-linear dimensionality reduction was performed using Uniform Manifold Approximation and Projection (UMAP) for visualization purposes. Clusters identified with the Louvain method were annotated based on their differentially expressed genes (computed using hurdle models for sparse single cell data from the MAST package) (Finak et al. 2015). Gene set enrichment analysis for molecular signatures was done using the method described by (Tirosh et al. 2016), as implemented in Seurat. In order to obtain a molecular signature of airway epithelial cell types that could be used to characterize iPSC-derived lung cells, we first analyzed a combination of two datasets of primary airway epithelial cells: (1) P0 HBECs in BEGM medium and (2) ALI cultures subsequently derived from these cells. We used the single cell pipeline above with one variation during the preprocessing: in addition to regressing the mitochondrial content, we also regressed out the "library" batch effect. Once we had annotated (based on the expression of known markers of basal, secretory and ciliated cells) the clusters obtained by the Louvain method (Figure S4B), we proceeded to compare each cell type against all others. In the cases where the cell type of interest was represented in more than one cluster, we combined such clusters before performing the comparison to ensure the purity of both query and background populations. This rationale is to ensure that the fold changes were not diluted by the presence of the same cell type in both the query and the background of each comparison, for example, when we grouped basal cell populations that were originally in different clusters belonging to HBEC samples and ALI samples, and to avoid noise by excluding clusters with poorly characterized or intermediate phenotypes.

All single cell visualizations were made with Seurat (heatmaps, UMAPs, violin plots).

The data discussed in this publication have been deposited in NCBI's Gene Expression Omnibus (Edgar et al. 2002) and are accessible through GEO Series accession number GSE142246 (https://www.ncbi.nlm.nih.gov/geo/query/acc.cgi?acc=GSE142246). Enter token “kjkbgqugdjmrdyd” into the box.

**Epithelial cell electrophysiological analysis**

Barrier function of the Air Liquid Interface Cultures (ALI) was measured using the Millicell ERS-2 Volt-ohm Meter (Millipore cat# MERS00002). The transepithelial electrical resistance (TEER) was measured at varying time points after the medium was removed from the apical chambers. All Transwells, including blanks (no cells), received 0.6ml of fresh PneumaCult-ALI medium in the basolateral chamber and 0.2ml in the apical chamber. Ohms (Ω) were measured by inserting the longer portion of the electrode into the lower chamber of the Transwell while the shorter end was kept in the apical chamber, and after readings were stabilized for at least six seconds. All measurements were performed in triplicate. TEER was obtained by calculating the average of the blank wells and subtracting them from the average of an individual Transwell’s 3 readings and multiplied by the Transwell surface area. TEER Ω. $cm^2$ = (Average(Ω)Transwell)-Average(Blank)* surface area ($cm^2$) of the Transwell.

Ussing chamber experiments were performed on EasyMount Ussing Chamber Systems at voltage clamp mode, and Acquire & Analyze software (Physiologic Instruments) was employed to record and analyze data. Briefly, Transwell inserts were mounted into chambers and bathed in low chloride Ringer's solution (1.2 mM NaCl, 140 mM Na-gluconate, 25 mM $NaHCO_3$, 3.33 mM $KH_2PO_4$, 0.83 mM $K_2HPO_4$, 1.2mM $CaCl_2$, 1.2 mM $MgCl_2$, 10 mM glucose) at apical and Ringer's solution (120 mM NaCl, 25 mM $NaHCO_3$, 3.33 mM KH2PO$_4$, 0.83 mM $K_2HPO_4$, 1.2mM $CaCl_2$, 1.2 mM $MgCl_2$, 10 mM glucose) at basolateral side of monolayer. After base line is stabilized, 100 µM amiloride (Sigma) applied to both sides of the chamber to carry out complete inhibition of ENaC (Epithelial sodium channel). Subsequently, 10 µM forskolin (Sigma) were administered to stimulate chloride current. At the end of experiments, 10 µM CFTR inhibitor-172 (Sigma) employed at the apical side to specifically inhibit CFTR function followed by 100 µM UTP at the apical side to assess the integrity of developed epithelium. For BU3 NGPT derived ALI, 1 µM VX-770, CFTR modulator, was treated subsequent to forskolin. The resulting change in short circuit current was calculated as ∆Isc. The data were expressed in mean±SD.

Ussing chamber analysis was also used to measure TEER at the beginning of each recording, prior to compound administration, in some cases. Briefly, a blank insert was mounted into the Ussing chamber and each side was incubated with Ringer solution as above, followed by compensation for voltage electrode asymmetry and fluid resistance. The blank insert was now replaced with the insert with cells, which was then voltage clamped and monitored for short-circuit current to calculate TEER based on Ohm's law.

**Limiting dilution and tracheosphere assays.**

On day 10 of directed differentiation, cells were treated with 5μg/ml polybrene and then separate wells were infected with pHAGE-EF1alphaL-eGFP-W, pHAGE-Ef1alphaL-TagBFP-W, or pHAGE-CMV-DsRED-W at an MOI=1 for 24 hr. On Day 20 of differentiation, cells were dissociated and GFP+, BFP+ and DsRed+ cells were sorted according to the same protocol described above. Equal numbers of GFP+, BFP+ and dsRED+ cells were combined and plated at 1000 cell/μL, 400 cells/μL and 200 cells/μL in 50ul droplets of 3D Matrigel in triplicate. On day 34, the Matrigel droplets were imaged and the number of GFP, BFP, dsRed or mixed colonies were manually counted. For the tracheosphere assay GFP+/TOM+/NGFR+ BU3 NGPT cells were sorted as above and plated at a density of 200cells/μL in basal cell medium. Once organoids had formed, ~ 10 days, triplicate wells were either continued in basal cell medium or changed to PneumaCult-ALI for an additional 10 days.

**Mucus cell metaplasia model**

BU3 NGPT iBSC-derived ALI culture was prepared as described above. Ten days after air-lifted, cells were treated with 10ng/ml IL-13 and cultured for 10 days prior to the assessments. Mucus cell metaplasia was assessed by immunofluorescence staining for the increment of MUC5AC expressing cells or gene expression changes via qRT-PCR. Quantification of MUC5AC+ cells was performed using ImageJ software by manually counting MUC5AC+ cells in 3 randomly selected fields per well (n=3 wells per condition).

**Reverse Transcriptase Quantitative RT Polymerase Chain Reaction (qRT-PCR)**

RNA was extracted by first lysing cells in QIAzol (Qiagen) and subsequently using the RNeasy Mini kit (Qiagen). Taqman Fast Universal PCR mastermix (Applied Biosystems) were used to reverse transcribe RNA into cDNA and analyzed during 40 cycles of real time PCR using Taqman probes (Applied Biosystems). Relative gene expression, normalized to 18S control, was calculated as fold change in 18S-normalized gene expression, over baseline, using the $2^{(-\Delta\Delta CT)}$ method. Baseline, defined as fold change = 1, was set to either undifferentiated iPSCs or HBEC-derived ALI as indicated in the figure legend. If undetected, a cycle number of 40 was assigned to allow fold change calculations.

**Transmission electron microscopy**

Electron microscopy was performed as previously described (Horani et al. 2013; Horani et al. 2012). In short, cultured airway epithelial cells collected from subjects with PCD, or iPSC derived cells were fixed using 2.5% glutaraldehyde + 2% paraformaldehyde in 0.15M cacodylate buffer at 37°C. Secondary fixation was performed in 1% $OsO_4$ / 1.5% potassium ferrocyanide, and then in 2% uranyl acetate. Post-staining with 0.1% lead citrate and 0.2% tannic acid was done for additional contrast enhancement. At least 10 cilia cross sections were evaluated per group.

**High speed video-microscopy**

Cilia beat frequency was analyzed live in at least 5 fields obtained from each preparation, using a high-speed video camera and processed with the Sisson-Ammons Video Analysis system (Amons Engineering, Mt Morris, MI) as described (Horani et al. 2012; Sisson et al. 2003).

**Statistics**

Statistical methods relevant to each figure are outlined in the accompanying figure legend. Unless otherwise indicated unpaired, two-tailed Student's *t*-tests were applied to two groups of n=3 or more samples, where each replicate ("n") represents either entirely separate differentiations from the pluripotent stem cell stage or replicates differentiated simultaneously and sorted into separate wells. A p-value <0.05 was considered to indicate a significant difference between groups.

| Gene | TaqMan Probe Number |
|---|---|
| *FOXJ1* | HS00230964 |
| *KRT5* | HS00361185 m1 |
| *KRT17* | HS00356958 m1 |
| *MUC5AC* | HS00873651 |
| *MUC5B* | HS00861595 m1 |
| *NGFR* | HS00609976 m1 |
| *SCGB3A2* | HS0036978 m1 |
| *SCGB1A1* | HS0036978 m1 |
| *TP63* | HS00978340 m1 |
| *SPEDF* | HS0017942 m1 |

**Table S1.** TaqMan Gene Expression Assay Information. Related to STAR Methods.

| Name | 5' to 3' |
|---|---|
| P63-15F | gttagcggatgctagggcaaatg |
| P63-5F | gtggcttcagcggctaata |
| P63-2R | agtgagagggaagcagaaatgaa |
| PGKrv | ccggtggatgtggaatgtgt |
| TKfw | tccgagacaatcgcgaacat |
| TKrv | accgtattggcaagtagccc |

**Table S2.** Primers used for characterization of the integrated reporter. Related to STAR Methods.
